# Non-canonical enhancers control gene expression and cell fate in human pluripotent stem cells

**DOI:** 10.1101/2025.06.01.657118

**Authors:** Yang Cao, Oleksandr Dovgusha, Ayyub Ebrahimi, Stephen Bevan, Lukas Widmer, Fereshteh Torabi, Julia Kurlovich, Ignacio Rodríguez-Polo, Soumen Khan, Manthan Patel, Madapura M Pradeepa, Ufuk Günesdogan, Stefan Schoenfelder

## Abstract

Enhancers are key gene regulatory elements that ensure the precise spatiotemporal execution of developmental gene expression programmes. However recent findings indicate that approaches to identify enhancers may not capture the full repertoire of active enhancers in mammalian genomes. Here, we combine massively parallel enhancer assays with chromatin structure and transcriptome profiling to functionally annotate enhancers genome-wide in human induced pluripotent stem cells. We find that a substantial fraction (∼40%) of accessible chromatin regions with enhancer function lack key features associated with active enhancers, including the active enhancer mark histone H3 lysine 27 acetylation and enhancer-associated RNAs. Perturbation of this class of non-canonical enhancers by CRISPR-mediated epigenome editing results in decreased levels of target gene expression and, in one instance, loss of pluripotent stem cell characteristics. Collectively, our data demonstrate enhancer activity for a class of gene regulatory elements that had until now only been associated with a neutral or inactive status, challenging current models of enhancer function.

## INTRODUCTION

Developmental gene expression programmes are controlled by transcriptional enhancers^1,2^, often over large genomic distances from their target genes^3,4^. The human genome is estimated to harbour up to one million mostly tissue-specific enhancers, vastly outnumbering genes^5^. Highlighting their importance in development, aberrant enhancer function or ectopic enhancer activation can lead to severe developmental malformations^6–8^. Further, thousands of disease-associated non-coding loci across dozens of disease indications carry hallmarks of enhancers^9^, and an increasing number of human diseases are associated with impaired enhancer activity^10,11^.

Enhancers also act as key regulators of the pluripotent stem cell (PSC) fate. Interfering with the function of enhancers that control key pluripotency genes results in the loss of mouse PSC properties^12^. Modulating the frequency of enhancer-promoter interactions is a key gene regulatory step during stem cell maintenance and differentiation into specialised cell types^13,14^. Epigenetic remodelling of enhancers has been shown to trigger reprogramming towards induced pluripotent stem cells (iPSCs)^15^, and targeting enhancers that control lineage-specifying genes can enhance the efficiency of differentiation from human PSCs into target cell types^16^. This illustrates the potential to experimentally target enhancers to direct cell fate decisions for stem cell-based applications in biomedicine.

Enhancers are predominantly located in the non-coding genome and are characterised by a high density of transcription factor binding sites. Enhancers were originally discovered through their ability to drive the expression of reporter genes^17^. However, most current genome-wide enhancer mapping strategies rely on transcription factor binding, such as EP300^18^, enrichment for specific post-translational histone modifications^19–21^, and/or the presence of enhancer-associated RNAs (eRNAs)^22^. In particular, specific combinations of post-translational histone tail modifications are widely used to classify enhancers into functional categories: inactive/neutral/primed (histone H3 lysine 4 mono-methylation (H3K4me1)), poised (H3K4me1 and histone H3 lysine 27 tri-methylation (H3K27me3)) and active (H3K4me1 and histone H3 lysine 27 acetylation (H3K27ac))^19–21^. Among these, H3K27ac is generally accepted as an active enhancer mark. However, recent findings suggest that not all, and possibly only a minority of, H3K27ac-enriched regions in the non-coding genome function as active enhancers. Only 12% of putative enhancers marked with H3K27ac in human PSCs function as transcriptional enhancers in a reporter gene assay^23^. Importantly, deletion of selected enhancers with reporter gene activity, but not of those without, affects the transcription levels of target genes, irrespective of H3K27ac enrichment^23^. Further, a substantial fraction (∼50%) of H3K27ac-positive and EP300-bound regions do not show enhancer activity *in vivo*^24–27^. Notably, H3K27ac is dispensable for enhancer activity in mouse embryonic stem cells (ESCs)^28^. Conversely, genomic regions devoid of H3K27ac have been shown to act as enhancers. In mouse ESCs, non-canonical enhancers have been identified that lack H3K27ac but instead are enriched for acetylation in the histone H3 core domain^29^. Studies dissecting gene control mechanisms at selected genomic loci in mouse (*Zfp42*, *Nanog*, *Rpp25*, *Tdgf1*) and human (*POU5F1*) PSCs have uncovered functional gene regulatory elements that lack canonical enhancer features^30,31^. Further, around one quarter of enhancers that are active *in vivo* in the mouse developmental regulator loci (*Gli3* and *Smad4/Smad6*) do not show canonical enhancer chromatin signatures in the corresponding tissues^26^. Together, these studies challenge the concept of H3K27ac as a universal mark of active enhancers, and raise the possibility that enhancer-like non-coding gene control elements exist in mammalian genomes which lack canonical enhancer chromatin signatures and thus evade detection by conventional chromatin profiling approaches. However, this has not been systematically addressed in a genome-scale manner in human PSCs.

Here, we combine a range of functional genomics methods including self-transcribing active regulatory region sequencing (STARR-seq^32^), ATAC-seq^33^, gene expression profiling by RNA-seq and transient transcriptome sequencing (TT-seq)^34^, CUT&Tag^35^ for post-translational histone modifications and Promoter Capture Hi-C (PCHi-C)^36,37^ to functionally annotate enhancers in human iPSCs (hiPSCs) genome-wide. Unexpectedly, we find that a substantial proportion of genomic regions with enhancer function in hiPSCs are devoid of the canonical active enhancer mark H3K27ac and have only low levels of eRNAs, thus escaping detection by conventional enhancer mapping approaches. We identify naturally occurring genetic variants in hiPSC cell lines from different donors at these putative non-canonical enhancers that impair their activity, which occur concomitantly with changes in chromatin accessibility and nearby gene expression. This establishes a link between non-canonical enhancer genetic variation, chromatin structure, enhancer activity and gene transcription. Further, we show that perturbation of putative non-canonical enhancer function at their endogenous chromatin loci by CRISPR interference leads to a reduction in the expression levels of target genes, demonstrating enhancer-like gene regulatory potential. In particular, we identified a non-canonical enhancer without active enhancer features that is required for cell survival and maintenance of pluripotency. Finally, non-coding putative regulatory elements with similar features also exist in mouse PSCs, and in human and *Drosophila* cell lines. Taken together, our results challenge the concept of H3K27ac as universal active enhancer mark, and define a novel class of non-canonical enhancers in human PSCs, suggesting that eukaryotic genomes may harbour tens of thousands of hitherto unknown enhancers that actively contribute to the control of developmental gene expression programmes.

## RESULTS

### Functional enhancer annotation in human induced pluripotent stem cells

To functionally annotate enhancers in hiPSCs, we performed genome-wide STARR-seq^38^. To this end, genomic DNA was sonicated and cloned into a STARR-vector downstream of a minimal promoter that is inducible by human enhancers^39^, and upstream of the 3’-untranslated region of a reporter gene. Sequencing of the input genome-wide STARR-seq library revealed 89% coverage of the mappable human genome, with a mean coverage of 5.9x. After transfecting the STARR-seq library into hiPSCs, we determined enhancer activity by RNA-seq. We generated three biological STARR-seq replicates from a hiPSC line generated by the HipSci consortium^40^ which we specifically selected for its representative pluripotency and novelty score (**Supplementary Figure 1A**)^41^, and absence of copy number variations (**Supplementary Figure 1B**). To our knowledge this is the first genome-wide functional enhancer atlas in a human non-cancer cell line. The three hiPSC STARR-seq replicates showed high correlation (**Supplementary Figure 1C, D**), allowing us to pool sequence reads for subsequent analyses.

Using a bespoke algorithm to correct for potential biases in STARR-seq data^42^, we identified 57,784 peaks in hiPSCs, 38,847 (67.3 %) of which were putative enhancer candidates as they were located in intronic and intergenic regions of the human genome. To correlate STARR-seq activity with chromatin accessibility at the endogenous enhancer loci, we generated omni-ATAC-seq^43^ libraries from hiPSCs. We identified 60,605 ATAC-seq peaks, 15.9% (9618) of which overlapped with STARR-seq enhancers. In line with the finding that PSCs are incapable of inducing an interferon type I response^44^, only 85 of the accessible hiPSC enhancers (0.9%) overlapped with a previously defined set of 1517 interferon-related enhancers that are induced by plasmid transfection^39^; these enhancers were excluded from further analyses. In addition, we omitted 29 putative enhancers that were enriched for the heterochromatin-associated mark tri-methylation of H3 lysine 9 (H3K9me3) at their endogenous chromatin loci.

We note that a major fraction of STARR-seq-positive genomic regions is located in inaccessible chromatin (**Figure 1A-C; Supplementary Table 1**), in agreement with previous studies^32,42,45,46^. Although inaccessible genomic regions can possess gene regulatory functions^47^ and have been shown to function as enhancers in some cases^26^, we conclude that the majority of STARR-seq+/ATAC-seq-enhancers most likely represent genomic sequences that have the ability to function as enhancers in episomal reporter assays, but whose activity is repressed by the endogenous chromatin context in hiPSCs. We here refer to this enhancer class as chromatin-repressed enhancers.

**Figure 1:**
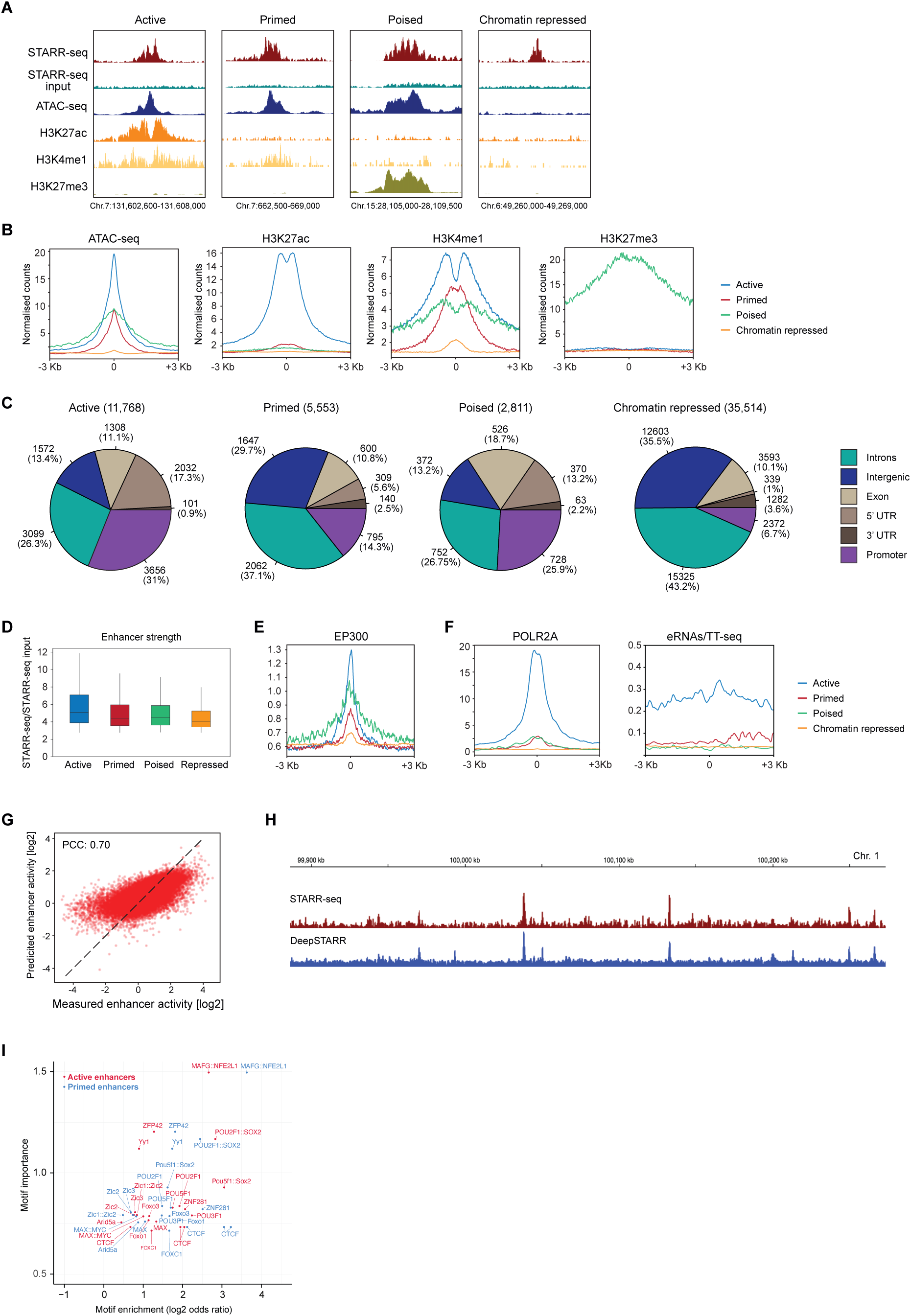
Identification and classification of putative enhancers in hiPSCs. (A) Coverage tracks of STARR-seq, ATAC-seq, ChIP-seq (H3K27ac, H3K4me1, H3K27me3) data in hiPSCs. Putative enhancers were mapped based on STARR-seq signal enrichment (top) over background control (STARR-seq input). Active, primed and poised enhancers are within accessible chromatin (ATAC-seq enrichment) and enriched for H3K4me1 and H3K27ac (active), H3K4me1 (primed), or H3K4me1 and H3K27me3 (poised). Chromatin-repressed enhancers are devoid of ATAC-seq signal and any of the analysed histone marks. Chr = chromosome. (B) Profile plots of ATAC-seq and ChIP-seq (H3K27ac, H3K4me1, H3K27me3) normalised counts at enhancers +/- 3kb of STARR-seq peak centres. (C) Numbers (and % of total) of enhancers within indicated genomic features. (D) Box plot showing enhancer strength, defined as STARR-seq signal over STARR-seq input control (background). (E) Profile plot of ChIP-seq EP300 normalised counts at enhancers +/- 3kb of STARR-seq peak centres. Colour legend shown in (F). (F) Profile plots of ChIP-seq RNA Polymerase II (POLR2A, left) and TT-seq (right) normalised counts at enhancers +/- 3kb of STARR-seq peak centres. (G) Pearson correlation (PCC) of DeepSTARR predicted versus observed enhancer activity genome-wide. (H) Example coverage tracks of STARR-seq data (red) in comparison to DeepSTARR predicted (blue) profiles. (I) Scatter plot showing motif enrichment (x-axis) and DeepSTARR-predicted global importance (y-axis) at active enhancers (red) and primed enhancers (blue). Note, only a subset of identified motifs is shown.

### Epigenetic modifications and transcription factor occupancy at hiPSC enhancers

Based on enrichment of post-translational histone modifications H3K4me1, H3K27ac, and H3K27me3, we classified STARR-seq+/ATAC-seq+ intergenic and intronic regions into three categories: 4671 active enhancers (enriched for H3K4me1 and H3K27ac), 3709 primed enhancers (enriched for H3K4me1), and 1124 poised enhancers (enriched for H3K4me1 and H3K27me3; **Figure 1A-C; Supplementary Tables 2-4**). As expected, active enhancers can be distinguished from the other enhancer classes by their higher STARR-seq activity (**Figure 1D**). Further, EP300 and RNA polymerase II occupancy were higher at active enhancers compared to primed and poised enhancers, concomitant with enhanced levels of eRNA transcription (**Figure 1E, F**). Collectively, our results establish the first genome-scale functional annotation of enhancers in human PSCs.

To identify enhancer sequence motifs, we used our STARR-seq data to train an enhancer syntax deep-learning model (DeepSTARR^48^). The predicted and observed enhancer activity profiles were highly similar genome-wide (Pearson correlation coefficient (PCC) = 0.70; **Figure 1G, H**), closely matching the reported predictive performance of DeepSTARR for *Drosophila* enhancers^48^. Next, we used the trained DeepSTARR model to predict the importance^48^ of enriched transcription factor motifs for active and primed enhancers. We found enrichment of various motifs of transcription factors involved in the control of pluripotency in both enhancer classes including POU5F1(OCT4), POU5F1::SOX2, ZFP281, YY1, and ZFP42(REX1) (**Figure 1I, Supplementary Figure 1E-H**). In addition, binding motifs for ZIC2 and ZIC3 were enriched, which are required to maintain primed pluripotency^49^. Taken together, the sequence motifs in active and primed enhancers are consistent with an important function in the maintenance of the pluripotent state.

### Long-range chromatin contacts and target genes of human pluripotent enhancers

Enhancers can control genes over large genomic distances, in some cases skipping over more proximally located genes to interact with their target genes^4^. To capture regulatory chromatin contacts between enhancers and all human gene promoters, we carried out PCHi-C^36,37^. To detect both mid- and long-range 3D chromatin interactions between enhancers and promoters, we generated PCHi-C maps using two different restriction enzymes (DpnII for higher resolution and mid-range interactions and HindIII for long-range interactions; **Supplementary Figure 2A; Supplementary Tables 5, 6**). Using HiCUP^50^, we confirmed that all PCHi-C libraries had a high percentage of valid reads (78.1-98%) and *cis* reads representing interactions on the same chromosome (71.6-84.6%; **Supplementary Figure 2B**), indicative of high quality chromatin interaction libraries. The individual PCHi-C libraries were sequenced to a depth of between 84.3 million and 170.6 million unique promoter-centered reads (**Supplementary Figure 2C**), totalling over 1.95 billion promoter interaction reads. CHiCAGO^51^ interaction calling uncovered 534,042 statistically significant promoter contacts (242,343 in HindIII PCHi-C libraries; 291,699 in DpnII PCHi-C libraries), representing the most comprehensive pan-human iPSC 3D promoter interactome atlas to date.

We next determined which gene promoters contact pluripotent STARR-seq enhancers (**Figure 2A**). This uncovered 2339, 1127, and 1079 putative target genes of active, primed and poised enhancers, respectively. Notably, we found that a major fraction of STARR-seq enhancers do not engage in promoter contacts, indicating that the promoter-enhancer contacts are either too dynamic to be captured, or that these enhancers may regulate their target genes without physical association in 3D nuclear space, as has been described for individual cases of ‘contactless’ enhancers^52,53^. As expected, genes interacting with active enhancers showed the highest expression level (**Figure 2B; Supplementary Figure 2D**). Assigning putative enhancers to their target gene based on genomic distance (+/-100kb) revealed the same trend (**Figure 2C**). Notably, genes interacting with, or located in proximity to, primed enhancers are more highly expressed than genes interacting with poised enhancers (**Figure 2B, C; Supplementary Figure 2D**), which have been shown to interact with their silenced target genes in pluripotent stem cells^54^.

**Figure 2:**
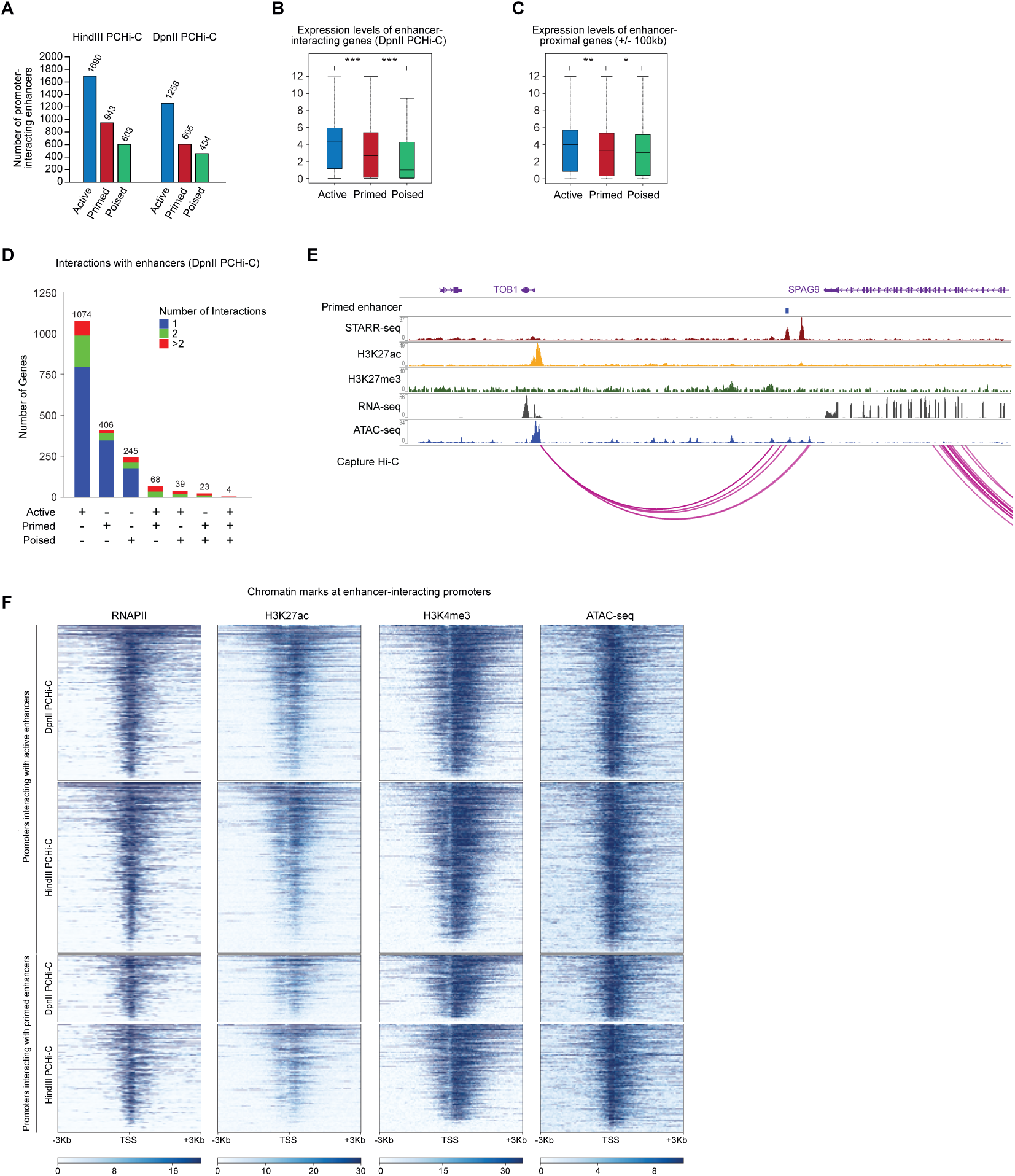
Long-range chromatin interactions of hiPSC enhancers. (A) Number of interactions of promoters with active, primed, or poised enhancers identified using HindII PCHi-C (left) and DpnII PCHi-C (right). (B) Expression levels (RNA-seq, log2 (normalised counts +1)) of enhancer interacting-genes. Interactions were mapped using DpnII PCHi-C data. (C) Expression levels (RNA-seq, log2 (normalised counts +1)) of enhancer-proximal genes (+/- 100kb). (D) Number of genes interacting with one, two or more than two active, primed and/or poised enhancers, respectively. Interactions were mapped using DpnII PCHi-C data. (E) Coverage tracks of genome-wide data on chromosome 17 showing a representative example of a primed enhancer, which interacts with an expressed gene (*TOB1*). Capture Hi-C corresponds to DpnII PCHi-C data. (F) Heatmap showing H3K27ac and H3K4me3 enrichment (ChIP-seq), and ATAC-seq signal, at promoters interacting with active or primed enhancers +/- 3kb of transcription start sites (TSSs) in primed hiPSCs. Interactions were mapped using HindIII PCHi-C or DpnII PCHi-C data.

Genes can in some cases receive regulatory inputs from more than one enhancer. To investigate more complex enhancer-promoter 3D networks, we separated genes by both the number and activity status of their interacting enhancer(s). This revealed that gene promoters preferentially interact with either active, primed or poised enhancers, whereas they engage with multiple enhancers harbouring different histone marks much less frequently (**Figure 2D, E; Supplementary Figure 2E-G**). This indicates that active, primed and poised 3D enhancer-promoter networks are largely spatially segregated in the nucleus of human PSCs, possibly contributing to a coordinated control of target gene expression levels. Finally, we determined chromatin accessibility (ATAC-seq), active chromatin marks (H3K4me3 and H3K27ac) and RNA polymerase II occupancy at the promoter regions of enhancer-interacting genes. Despite clearly distinct gene expression levels (**Figure 2B, C; Supplementary Figure 2D**), we detected only minor differences in RNAPII occupancy and H3K27ac enrichment at promoters interacting with active enhancers compared to promoters interacting with primed enhancers, and the H3K4me3 and ATAC-seq profiles were nearly indistinguishable (**Figure 2E**). Thus, chromatin accessibility, histone marks (**Figure 1B**) and RNAPII occupancy (**Figure 1F**) at enhancers appear to correlate more strongly with gene expression levels than the corresponding marks at the enhancer-interacting gene promoters.

### Chromatin mark dynamics at hiPSC enhancers during cell lineage specification

Primed enhancers are thought to exist in a neutral and inactive state, poised for activation upon acquisition of the active histone mark H3K27ac during cell lineage diversification. To test this prediction for the primed hiPSC enhancers (STARR-seq+ ATAC-seq+ H3K27ac-), we used previously published epigenome profiling data from cell types that represent key lineages in the human embryo: trophoblast-like cells (TBL), mesendoderm (ME), neural progenitor cells (NPCs) and mesenchymal stem cells (MSCs)^55^. We found that 19.1% of candidate primed enhancers (707/3709) gained H3K27ac specifically in one of the four lineages (‘lineage-restricted’; MES 9.4%; TBL 3.4%; NPC 3.9%; MSC 2.3%), whereas 13.8% of candidate primed enhancers (511/3709) gained H3K27ac in all four lineages (‘multi-lineage’; **Figure 3A**). However, the majority (2491/3709; 67.1%) of primed enhancers failed to acquire the active enhancer mark H3K27ac in any of the cell types interrogated (**Figure 3A, B**), comparable to poised and chromatin-repressed enhancers (**Figure 3B; Supplementary Figure 3A, B**).

**Figure 3:**
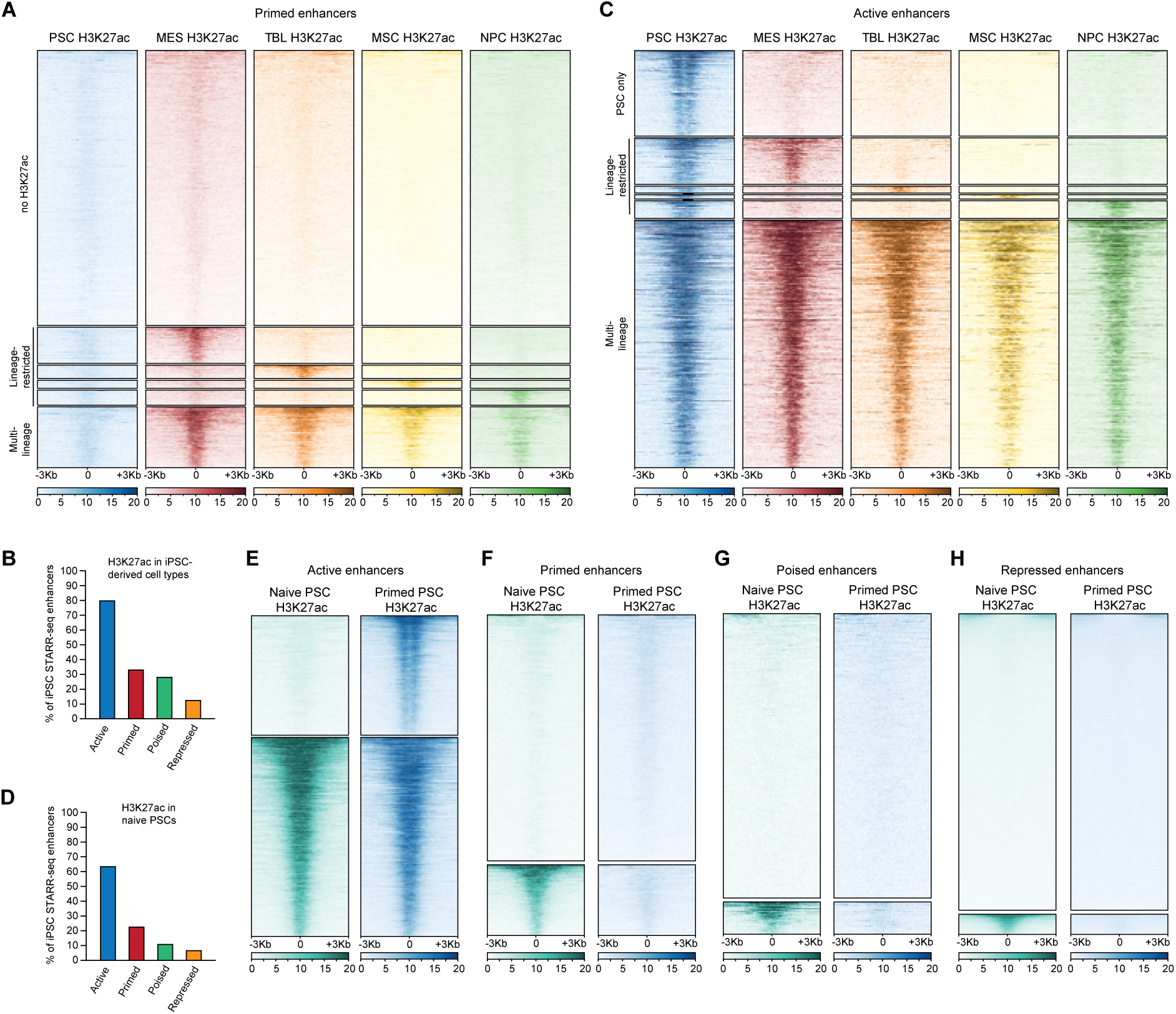
The majority of primed enhancers do not gain H3K27ac during differentiation. (A) Heatmap showing H3K27ac enrichment (ChIP-seq) at primed enhancers +/- 3kb of STARR-seq peak centres in primed human pluripotent stem cells (PSC), mesendoderm (MES), trophoblast-like cells (TBL), mesenchymal stem cells (MSC), and neuronal progenitor cells (NPC). The majority of enhancers are not enriched for H3K27ac in any of the analysed cell types (no H3K27ac). Subsets are enriched for H3K27ac in one (Lineage-restricted) or multiple differentiated cell types (Multi-lineage). (B) Proportion (in %) of enhancers mapped in hiPSCs that maintain or gain H3K27ac in one or multiple differentiated cell types. (C) Heatmap showing H3K27ac enrichment (ChIP-seq) at active enhancers +/- 3kb of STARR-seq peak centres in PSC, MES, TBL, MSC, and NPC. Subsets of enhancers are enriched for H3K27ac only in hiPSCs (PSC only), or in hiPSCs and one (Lineage-restricted) or multiple cell types (Multi-lineage). (D) Proportion (in %) of enhancers mapped in primed hiPSCs that are enriched for H3K27ac in naïve PSCs. (E) – (H) Heatmap showing H3K27ac enrichment (ChIP-seq) at active (E), primed (F), poised (G), and repressed (H) enhancers (mapped in primed hiPSCs) +/- 3kb of STARR-seq peak centres in naïve and primed PSCs.

In contrast, the majority of active enhancers (STARR-seq+ ATAC-seq+ H3K27ac+), maintained H3K27ac in all lineages (2771/4671; 59.3%), whereas 20.1% of active enhancers maintained H3K27ac selectively in one lineage (‘lineage-restricted’; 938/4671; MES 12.1%; TBL 1.9%; NPC 4.8%; MSC 1.3%). Notably, only 20.6% of active enhancers (962 enhancers) harbour H3K27ac exclusively in hiPSCs (**Figure 3B, C**). We next tested whether primed enhancers carry active enhancer marks in naïve hPSCs, which represent an earlier stage of human development. Using published H3K27ac data^56^, we established that the majority of active hiPSC enhancers are marked with H3K27ac already in naïve hPSCs (**Figure 3D, E**). In contrast, the majority of primed, poised and chromatin repressed enhancers are not enriched for H3K27ac in naïve hPSCs (**Figure 3D, F-H**). Thus, we conclude that most primed enhancers in hiPSC are not occupied by H3K27ac in either developmental progenitor states or major cell lineages derived from hPSCs. This observation is consistent with enhancer priming for activation in cell types or states that we have not interrogated in this study, including at later developmental stages. Alternatively, primed enhancers may have yet to be identified functions in human PSCs.

### Sequence determinants of human pluripotent enhancers

Mutations in transcription factor binding sites within enhancers can lead to severe developmental malformations^57^, and a growing number of diseases and disease susceptibilities have been linked to aberrant enhancer function caused by genetic variation^9,11^. Deciphering how enhancer variants affect developmental gene expression programmes and shape complex human traits is therefore a central goal of the ‘variant-to-function’ challenge. However, our understanding of how naturally occurring genetic variation controls enhancer function in human PSCs is limited.

To identify functional enhancer variants, we pursued a two-pronged approach. First, we interrogated our STARR-seq data for heterozygous enhancer sequence variants that result in inter-allelic differences in enhancer activity. This approach revealed 15 and 16 functional (differences between major and minor enhancer alleles two-fold or more) enhancer variants in active and primed enhancers, respectively (**Figure 4A, B**). Next, as the absence of a haplotype-phased genome assembly for the hiPSC line used for our STARR-seq experiments prevented us from unambiguously assessing the effect of inter-allelic enhancer variants on the expression level of target genes, we focussed on homozygous enhancer SNPs that differ between individual hiPSC lines. As a proof of concept, we tested genetic variants for one primed and one chromatin-repressed enhancer from our STARR-seq data in a luciferase enhancer assay (**Figure 4C**), using established negative and positive controls from the *FGFR1*^23^ and *POU5F1*^31^ loci, respectively (**Supplementary Figure 4A, B**). We generated matching ATAC-seq and RNA-seq data from a hiPSC line in which these enhancers have opposing activities, i.e. enhancer 1 (Enh1) is a primed enhancer in the hiPSC line derived from donor 1, but a chromatin-repressed enhancer in the hiPSC line derived from donor 2 (**Figure 4D**), whereas enhancer 2 (Enh2) is a chromatin-repressed enhancer in the hiPSC line derived from donor 1, but a primed enhancer in the hiPSC line derived from donor 2 (**Figure 4E**). For both enhancers, we detected pronounced inter-individual differences in enhancer activity (**Figure 4C**) which correlated positively with ATAC-seq peak strength and the expression levels for specific genes at the endogenous loci (*EEIG2* for Enh1; *ABHD4* for Enh2; **Figure 4D, E**). Further, the relative enhancer strengths were correctly predicted by DeepSTARR^48^ (**Supplementary Figure 4C**). These results indicate that genetic variants in active, primed and chromatin-repressed enhancers have the potential to shape inter-individual differences in gene expression programmes between hiPSC lines derived from different donors.

**Figure 4:**
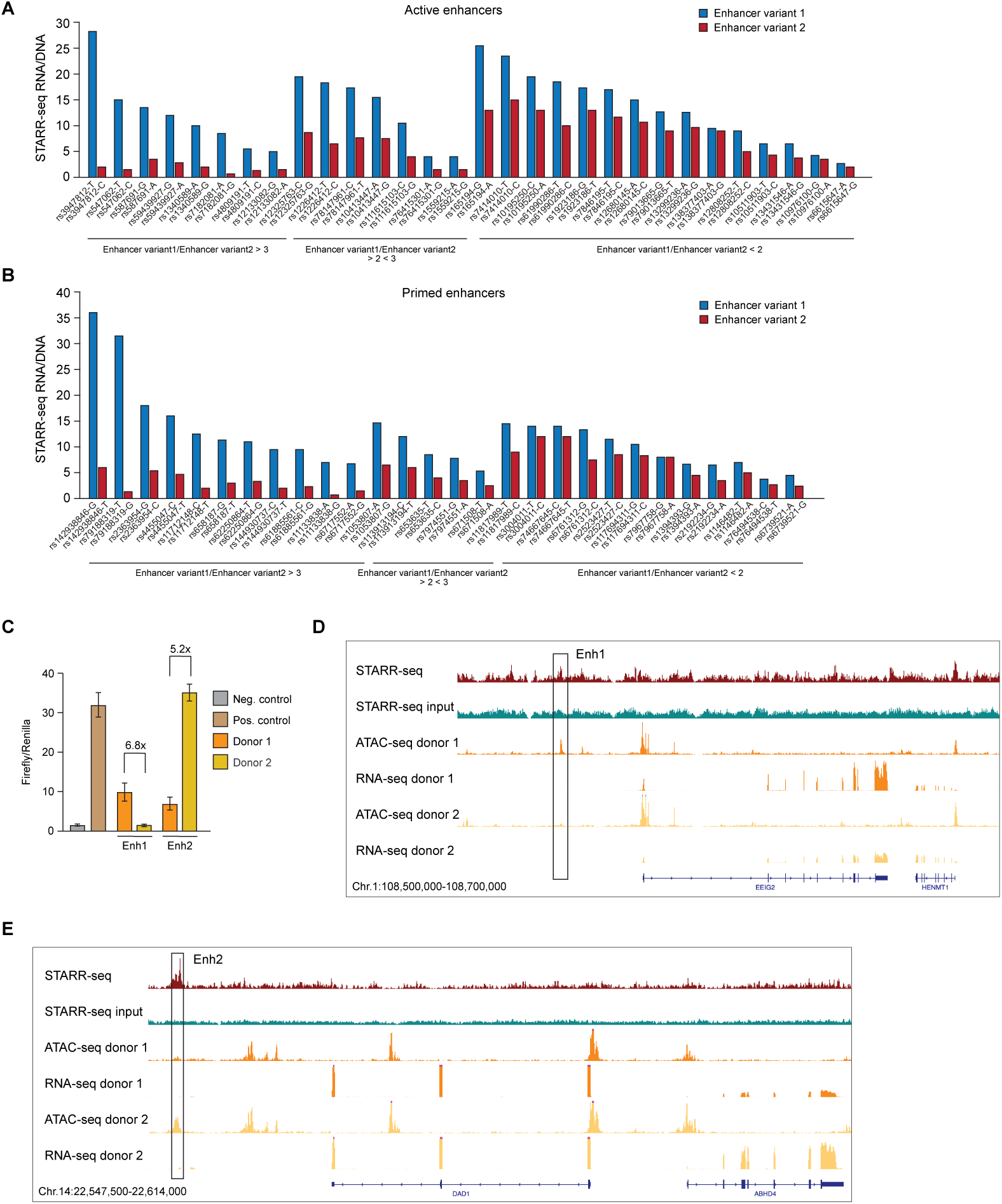
Sequence variation impairs the activity of active and primed enhancers. (A) Enhancer activity measured by STARR-seq at active enhancers with inter-allelic sequence variation. (B) Enhancer activity measured by STARR-seq at primed enhancers with inter-allelic sequence variation. (C) Luciferase reporter assay for genetic variants in two hiPSC STARR-seq enhancers (Enh1 and Enh2). Enhancer sequences were derived from HipSci consortium whole genome sequencing data (https://www.hipsci.org/#/) of two independent hiPSC lines from Donor 1 and Donor 2. Negative control: element from the *FGFR1* superenhancer that lacks enhancer activity (Barakat et al., 2018); Positive control: distal enhancer of *POU5F1*. (D) Coverage tracks of STARR-seq (donor 1) and ATAC-seq and RNA-seq data from two independent hiPSC lines (donor 1 and donor 2) for Enhancer 1 (Enh1; see Fig. 4C). (E) Coverage tracks of STARR-seq (donor 1) and ATAC-seq and RNA-seq data from two independent hiPSC lines (donor 1 and donor 2) for Enhancer 2 (Enh2; see Fig. 4C).

### CRISPR enhancer perturbation reveals primed enhancer activity in human PSCs

Given that (i) the majority of primed STARR-seq enhancers do not acquire active enhancer signatures in key PSC-derived lineages (**Figure 3B, C**), (ii) sequence variants affect primed enhancer strength in hPSCs (**Figure 4C-E**), and iii) primed enhancers interact with genes expressed at moderate levels (**Figure 2B; Supplementary Figure 2D**), it is possible that primed enhancers control gene expression in hiPSCs, contradicting current models of primed enhancer function. To test this hypothesis directly, we used a doxycycline-inducible CRISPR inactivation/activation system^16^ to modulate enhancer function within the endogenous chromatin contexts. Targeting the KRAB transcriptional repressor domain to enhancers (CRISPR interference, CRISPRi), a widely used method to interrogate enhancer function^58–60^, resulted in transcriptional downregulation of the putative primed enhancer target genes *APLN*, *FAM102B*, *XPNPEP2*, *LAGE3* and *SLC13A4*, at levels comparable to the active enhancer target gene *SFRP2*^23^, which we used as a positive control (**Figure 5A**). Conversely, targeting the transcriptional activator VP64 (CRISPR activation; CRISPRa) to primed enhancers resulted in transcriptional upregulation of target genes *APLN* and *XPNPEP2* (**Figure 5B**). These results demonstrate that interfering with primed enhancer activity results in changes in target gene expression in hPSCs, challenging the current model for primed enhancer function which predicts that they become active only at later developmental stages in more specialised cell types.

**Figure 5:**
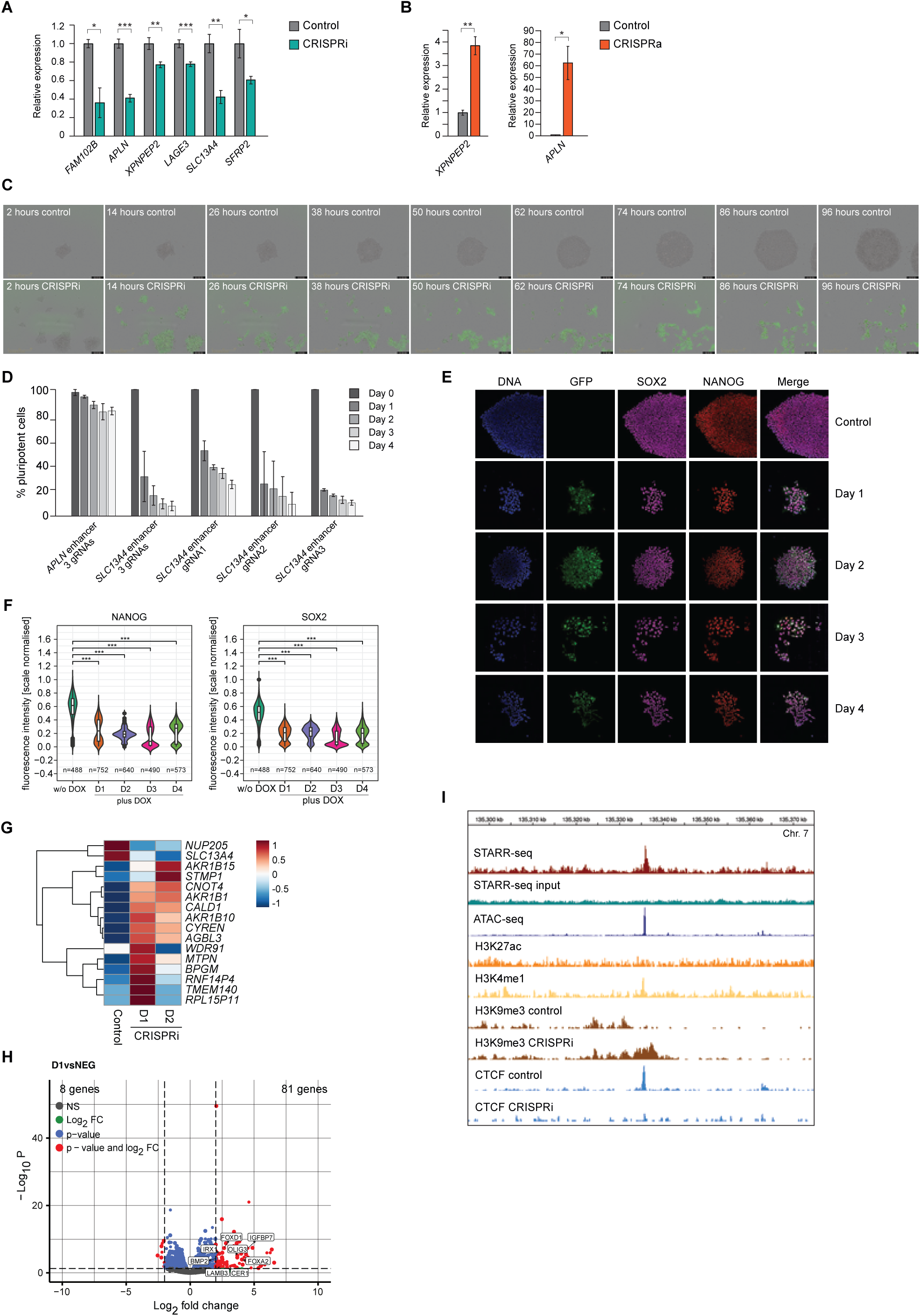
Repression of primed enhancers reduces target gene expression. (A) Quantitative RT-PCR for indicated genes before (Control) or after DOX-induced expression of dCas9-KRAB (CRISPRi) and gRNA-mediated targeting to primed enhancers in hiPSCs. 2^-ΔΔCt^ ± s.d.; normalisation using GAPDH (housekeeping gene) and uninduced hiPSCs as a reference (= 1). (B) Quantitative RT-PCR for indicated genes before (Control) or after DOX-induced expression of dCas9-VP16 (CRISPRa) and gRNA-mediated targeting to primed enhancers in hiPSCs. 2^-ΔΔCt^ ± s.d.; normalisation using GAPDH (housekeeping gene) and uninduced hiPSCs as a reference (= 1). (C) Live-cell imaging of hiPSCs without (Control) or with (CRISPRi) the addition of DOX and TMP to induce the expression of dCas9-KRAB, marked by GFP expression (green). Constitutive expression of three gRNAs targeting the primed enhancer of *SLC13A4*. (D) Proportion of remaining cells (in %) before (Day 0) or after addition of DOX and TMP (Day1-4) for dCas9-KRAB expression during live-cell imaging. Three (3 gRNAs) or individual gRNAs (gRNA1-3) for targeting the enhancer of *APLN* or *SLC13A4* were constitutively expressed. (E) Immunofluorescence images of hiPSC colonies before (Control) or after addition of DOX and TMP (Day 1-4) for dCas9-KRAB expression, marked by GFP expression. Three gRNAs targeting the enhancer of S*LC13A4* were expressed constitutively. Samples were stained for DNA, SOX2, and NANOG. (F) Quantification of fluorescence intensity of SOX2 and NANOG in images shown in (E). (G) Heatmap shows relative expression levels of genes +/- 1Mb of the *SLC13A4* locus based on RNA-seq data for hiPSCs after the addition of DOX and TMP for 1 day (D1) or 2 days (D2) to induce dCas9-KRAB with constitutive expression of gRNAs targeting the enhancer of *SLC13A4* in comparison to control hiPSCs. (H) Volcano plot shows differentially expressed genes of RNA-seq data for hiPSCs after the addition of DOX and TMP for 1 day to induce dCas9-KRAB with constitutive expression of gRNAs targeting the enhancer of *SLC13A4* in comparison to control hiPSCs. (I) Coverage tracks of STARR-seq, ATAC-seq, ChIP-seq (H3K27ac, H3K4me1) showing the *SLC13A4* enhancer region in hiPSCs, and H3K9me3 and CTCF CUT&Tag tracks showing the same *SLC13A4* enhancer genomic region in control and enhancer CRISPRi conditions. Chr = chromosome.

### A CTCF-bound primed enhancer is required for human pluripotency

Notably, CRISPRi targeting of one primed enhancer resulted not only in *SLC13A4* transcriptional downregulation (**Figure 5A**), but also in pronounced changes of hiPSC colony morphology and substantial cell death, as revealed by live-cell imaging over 4 days (**Figure 5C; Supplementary Movie 1**). In contrast, in non-CRISPRi-induced control hiPSCs, colony formation, hiPSC morphology and cell survival were unaffected (**Figure 5C; Supplementary Movie 2**). Additionally, CRISPRi targeting of the primed enhancer controlling *APLN* had only minor effects on cell viability (**Figure 5D**) and morphology (**Supplementary Figure 5A; Supplementary Movies 3, 4**). The changes in hiPSC colony morphology could be recapitulated with all three *SLC13A4* enhancer-targeting gRNAs used individually in CRISPRi, albeit to differing degrees (**Figure 5D; Supplementary Figure 5B-D**). Further, we recapitulated the changes in hiPSC morphology upon *SLC13A4* enhancer perturbation in an independent transfection using a different hiPSC line (**Supplementary Movies 5, 6**), demonstrating that neither donor genetic background nor the genomic integration sites of the dCas9-KRAB and gRNA plasmids can account for the observed phenotype. As an additional control, we detected no correlation between dCas9 induction levels and the severity of the induced phenotype, arguing against unspecific effects on cell fitness caused by dCas9-KRAB overexpression (**Supplementary Figure 5E**).

Although the remaining hiPSC colonies expressed the pluripotency markers SOX2 and NANOG after *SLC13A4* enhancer perturbation (**Figure 5E**), quantification of the immunofluorescence images revealed reduced protein levels for both pluripotency factors (**Figure 5F**).

To determine whether the *SLC13A4* enhancer controls additional target genes, we carried out RNA-seq in hiPSCs on day 1 and 2 after dCas9-KRAB induction. Within a 2Mb genomic window around the enhancer, only *NUP205*, a component of the nuclear pore complex, and *SLC13A4* showed consistent downregulation upon CRISPRi, whereas several other genes in the locus displayed moderate expression changes including mild transcriptional upregulation, and oscillations between up- and downregulation (**Figure 5G**). At the genome-scale level, we detected upregulation of several differentiation markers upon enhancer CRISPRi, including *FOXD1*, *FOXA2*, *IGFBP7* and *OLIG3* (**Figure 5H**). Gene ontology analyses of differentially expressed genes revealed an enrichment for tissues of mesodermal origin, consistent with aberrant expression of *FOXD1* and *FOXA2* (**Figure 5H; Supplementary Figure 5F**). Together, these findings are indicative of a partial loss of pluripotent stem cell features as a consequence of impaired primed *SLC13A4* enhancer function.

To control for possible CRISPRi off-target effects, we next carried out CUT&Tag against H3K9me3, the histone modification induced by dCas9-KRAB. We found that H3K9me3 spreading is limited to a region of ∼ 6 kb around the genomic site targeted by the gRNAs (**Figure 5I**), without detectable changes to H3K9me3 levels on any of the gene promoters or gene bodies in the *SLC13A4* enhancer locus. Consensus CTCF binding sites are amongst the most strongly enriched sequence motifs of primed hiPSC enhancers (**Figure 1I; Supplementary Figure 1E-H**). Notably, CUT&Tag revealed pronounced CTCF binding to the *SLC13A4* enhancer (**Figure 5I**). Upon CRISPRi, CTCF binding to the enhancer is markedly reduced (**Figure 5I**), indicating that CTCF occupancy may be required for appropriate enhancer function. Collectively, our data identify a hitherto uncharacterised gene control element as an essential component of the gene regulatory circuitry that maintains the pluripotent state of hiPSCs.

### Putative non-canonical enhancers are widespread in the human genome

Our results demonstrate that some primed enhancers can function as *bona fide* enhancers in human pluripotent stem cells, despite lacking the active enhancer mark H3K27ac. Similar enhancer activities have been reported for non-canonical enhancers in mouse ESCs and human K562 cells^29^, which are marked by acetylated lysine residues in the globular domain of histone H3, including lysine 122. We found that H3K122ac is not enriched at primed compared to active STARR-seq enhancers in hPSCs (**Figure 6A**), suggesting that the enhancers we uncover here are functionally distinct from mouse stem cell non-canonical enhancers. In contrast, we found that acetylation of histone H4 lysine 12 (H4K12ac) is moderately enriched at primed over active enhancers in hPSCs (**Figure 6B, C**). We next asked whether the putative novel enhancer class our data has revealed are only found in hPSCs. Re-analysing publicly available genome-wide STARR-seq datasets^32,42,45,46^, we found that putative non-canonical enhancers (H3K27ac-, ATAC-seq+, STARR-seq+) can also readily be identified in human cancer cell lines (LnCAP, K562, HepG2, HeLa), and to a lesser extent in mouse embryonic stem cells and *Drosophila* S2 cells (**Figure 6D**). This observation cannot be explained by the number of H3K27ac peaks called in the respective data sets (**Supplementary Figure 6A**). Further, the genomic locations of the putative non-canonical enhancers are consistent with their function as transcriptional enhancers (**Supplementary Figure 6B**). Although targeted enhancer perturbation experiments are currently lacking to directly test this hypothesis in cell types other than human PSCs, these findings suggest that non-canonical enhancers devoid of the active enhancer mark H3K27ac may be widespread in the human genome across a broad range of cell types.

**Figure 6:**
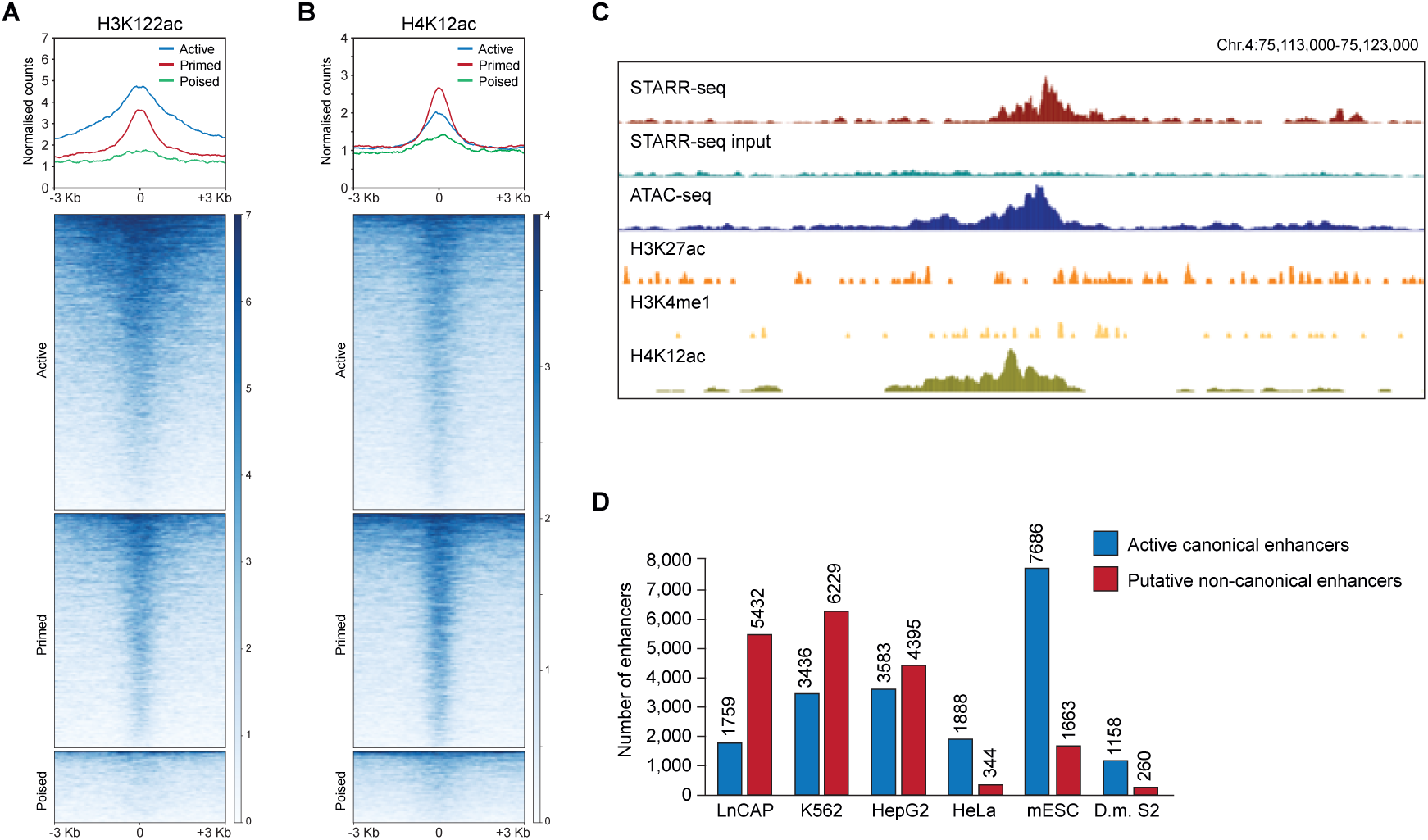
Putative non-canonical enhancers in Drosophila, mouse and human cells. (A) Profile plot of CUT&Tag H3K122ac at active, primed and poised enhancers +/- 3kb of STARR-seq peak centres. (B) Profile plot of CUT&Tag H4K12ac at active, primed and poised enhancers +/- 3kb of STARR-seq peak centres. (C) Coverage tracks of STARR-seq, ATAC-seq, ChIP-seq (H3K27ac, H3K4me1), and H4K12ac CUT&Tag, demonstrating an example of H4K12ac enrichment over the indicated primed enhancer. Chr = chromosome. (D) Number of active canonical and putative non-canonical enhancers in published STARR-seq datasets from LnCAP, K562, HepG2, HeLa, mouse ESC, and *Drosophila melanogaster* S2 cells.

## DISCUSSION

Here, we present the first genome-scale functional annotation of enhancers in human pluripotent stem cells, using a combination of massively parallel reporter gene assays and functional genomics profiling approaches, supported by live cell imaging and CRISPR epigenome editing.

Among our key findings, we show that enhancers lacking the active enhancer mark H3K27ac can function as *bona fide* enhancers in human pluripotent stem cells. This class of enhancers (ATAC-seq+, H3K4me1+, H3K27ac-) has previously been described as primed, neutral or inactive enhancers^19–21,61^. The current model of primed/neutral/inactive enhancer function predicts that this class of enhancers is transcriptionally inert and only controls target gene expression upon acquisition of the active mark H3K27ac in more specialised cell types. Our findings directly challenge this model, as we demonstrate that primed enhancer perturbation results in reduced transcriptional output from target genes in iPSCs. Underscoring the functional importance of this enhancer class for human pluripotency, we find that CRISPRi targeting of one primed enhancer results in impaired cell viability and loss of pluripotency features, including changes in stem cell colony morphology, reduced pluripotency marker expression levels and aberrant expression of differentiation markers. At the gene level, CRISPR inactivation of this enhancer leads to overall subtle expression changes of several genes in the locus, with only *NUP205* and *SLC13A4* displaying reduced expression. *NUP205* encodes an inner nucleoporin subunit of the nuclear pore complex, and mutations in *NUP205* have been shown to cause steroid-resistant nephrotic syndrome^62^ and abnormal cardiac left-right patterning^63^. *SLC13A4* encodes a transmembrane protein that controls the transport of sulfate which is essential for various metabolic pathways. Conditional knockout of *SLC13A4* in the placenta causes embryonic lethality, suggesting that the control of sulfate transport to the fetus is critical for development^64,65^. Further, *Slc13a4* haplo-insufficient mice show impaired postnatal brain development^66^. In hiPSCs, *SLC13A4* is expressed at relatively low levels, which at first glance makes it an unlikely candidate for a gene whose misexpression causes a pronounced effect on iPSC viability and morphology. However, we note that similarly lowly expressed genes such as *FANCB* and *FAM71F1* are essential for human pluripotency^67^. Thus, it is conceivable that subtle changes in the expression levels of *NUP205* and *SLC13A4* are causal for the impaired iPSC viability we report here. An alternative explanation is that the phenotype we observe is the combined result of a transcriptional imbalance across multiple genes in the locus upon enhancer perturbation. The precise mechanisms of action of this enhancer remains to be elucidated in future studies.

Here, we report the discovery of a class of non-canonical enhancers in hPSCs, which lack the two most commonly used markers to identify active enhancers: H3K27ac and enhancer RNAs. This raises questions as to how universally applicable these marks are for active enhancer discovery. Acetylation of H3K27 is mediated by the transcriptional co-activator and histone acetyltransferase CBP/EP300 neutralising the positive charge of histone H3. This is thought to lead to chromatin decondensation to increase accessibility for binding of transcription factors to activate enhancers^68^. Notably, however, the chromatin accessibility landscape is largely unaffected upon genome-wide hypoacetylation by p300/CBP inhibition^69^. Moreover, H3K27ac enrichment at enhancers does not necessarily correlate with their activity, as the majority of H3K27ac-enriched enhancers tested in human ESCs or in mouse embryos does not drive reporter gene expression^23,24,26,27^. Importantly, a mutation of the H3K27 residue (H3K27R) in flies or mouse ESCs does not interfere with enhancer activity and gene induction during differentiation, suggesting that the acetylation of H3K27 is dispensable for transcription^28,70^. The concomitant lack of H3K27 methylation in H3K27R mutant cells results in transcriptional derepression of PRC2 target genes, indicating that H3K27ac might serve as a ‘placeholder’ to prevent H3K27 methylation^28,70^. These studies suggest that H3K27ac may not be essential for enhancer activation, which could also be due to a degree of redundancy as other histone residues can also be acetylated. In mouse ESCs, active non-canonical enhancers are enriched for H3K122ac and H3K64ac^29^. Similarly, histone H2B acetylation was found at active enhancers in mouse ESCs or human cells^71^. The active non-canonical enhancers identified in this study lack H3K27ac but do show moderate enrichment for H3K122ac and H4K12ac, pointing to a more complex histone modification enhancer code (further discussed below).

Active canonical enhancers are bound by EP300/CBP^18^ and RNAPII, and are bi-directionally transcribed into non-coding eRNAs^22^. In contrast, our data suggests that non-canonical enhancers are bound by EP300/CBP but lack RNAPII occupancy and eRNA expression. This is consistent with the absence of H3K27ac at non-canonical enhancers, since the histone acetyltransferase activity of CBP is enhanced by the interaction with eRNAs^72^. The level of eRNA expression at active canonical enhancers is highly variable^73^. Functionally, they can promote target gene expression, chromatin looping and enhance transcription factor occupancy^22^. Thus, the lack of eRNA transcription at non-canonical enhancers may be related to the lower levels of target gene expression as compared to canonical enhancer target genes.

This study has limitations that should be acknowledged. First, STARR-seq does not measure the activity of enhancers in their endogenous chromatin environments, and a previous study has reported pronounced differences in enhancer activities measured through episomal versus chromatin-integration assays^74^. Here, we mitigate this limitation by integrating enhancer activity measured by STARR-seq with a range of complementary functional genomics assays that directly assess chromatin accessibility, posttranslational histone modifications and transcriptional activity at the endogenous enhancer loci. Nonetheless, the number of enhancers which we have interrogated directly by CRISPR perturbation is limited, and future studies will address on a more comprehensive scale how widespread genuine enhancer activity is among human iPSC primed enhancers. Second, we have determined enhancer activity across the human pluripotent genome using only one promoter in our STARR-seq approach, namely the origin of replication (ORI) core promoter. Even though ORI has been thoroughly tested and is established as a highly inducible promoter that responds to human enhancers across a range of human cell types^39^, *Drosophila* enhancers have been shown to exhibit marked specificity to different promoters^75^, and we cannot with certainty exclude the possibility that some human pluripotent enhancers may not be functionally compatible with the ORI promoter.

Notably, we found a large number of intronic and intergenic H3K27ac marked regions in pluripotent open chromatin (28,206/33,183; 85 %) that display no or very little STARR-seq enhancer activity, in line with a previous study^23^. The corresponding enhancer class (STARR^-^, ATAC^+^, H3K27ac^+^) in GP5d and HepG2 cells has recently been termed ‘chromatin-dependent’, and it has been suggested that these enhancers cannot be effectively detected using STARR-seq^76^. While this remains a plausible explanation for at least some enhancers, biochemical assays such as H3K27ac ChIP-seq (or H3K27ac CUT&Tag) profile a chromatin-associated mark that is correlated with enhancer activity, rather than directly demonstrating enhancer function. Notably, CRISPR deletion of H3K27ac+ STARR+ candidate enhancers resulted in reduced transcriptional output of target genes, whereas deletion of H3K27ac+ STARR-putative enhancers did not^23^. Thus, enhancer function cannot be inferred from chromatin signatures alone, and large-scale enhancer perturbation approaches will be required to determine how many H3K27ac marked putative enhancers possess genuine enhancer activity, and in which cellular contexts.

In sum, our data identify a novel class of non-canonical enhancers with essential functions in human pluripotent stem cells. Our findings, along with those of others^23,24,26–31,77^, indicate that the classification of enhancers into only three functional categories (active, primed and poised) based on their posttranslational histone modification signatures does not accurately reflect the complexity of enhancers and their associated gene regulatory landscapes in mammalian cells. We predict that combinations of posttranslational histone tail and core modifications^19–21,29,78^, including the recently discovered acetylation of histone H2B^71^, will dictate enhancer function and activity. Future studies will aim to refine how this ‘enhancer histone code’ translates signalling inputs into regulatory information relayed to target genes, and how cell type and developmental context dependencies shape the activities of the diverse enhancer functional subcategories on the transcriptional outputs of their target genes.

## METHODS

### Cell culture

hiPSC lines in this study were purchased from the Human Induced Pluripotent Stem Cells Initiative (https://www.hipsci.org). HPSI0215i-yoch_6 was used for STARR-seq, TT-seq and CUT&Tag; HPSI0215i-yoch_6 and HPSI1113i-qolg_1 were used for luciferase assays, ATAC-seq and RNA-seq; and HPSI0215i-yoch_6 and HPSI0114i-kolf_2 were used for CRISPR. All cell lines were cultured in TeSR-E8 media with supplement (Stemcell Technologies, 05990) on Vitronectin (Thermo Fisher Scientific, A14700) coated surfaces at 37°C under 5% CO_2_ and normal O_2_ levels. Media with supplement was refreshed every 24 hours. 10 µM Rho-associated protein Kinase (ROCK) inhibitor (Cell Guidance Systems, Y-27632) was added to the culture for the first day after revival. Cells were passaged at ratios ranging from 1/6 to 1/8 using 0.5 mM EDTA when they were approximately 70% confluent. For electroporation transfections in STARR-seq screening and luciferase assays, cells were cultured in TeSR-E8 media with supplement (Stemcell Technologies, 05990) and 10 µM ROCK inhibitor for 24 h before electroporation, and 6 h (STARR-seq candidates) or 24 h (STARR-seq controls and luciferase assay) after electroporation and recovery. After the first 24 h, cells in STARR-seq controls were cultured for additional 24 h without Rho-associated protein Kinase (ROCK) inhibitor. Prior to experiments, cells were harvested and detached using Accutase solution (Sigma-Aldrich, A6964). Cell number and viability were analysed using Countess Cell Counting Chamber Slides (Invitrogen, C10228) on the Countess II Automated Cell Counter (Invitrogen). Once revived, only the cells with no less than three passages and no more than ten passages were harvested for experiments.

### ATAC-seq

ATAC-seq libraries were generated as described^43^, with minor modifications. 50,000 cells were harvested with Accutase, and centrifuged at 500g for 5 min at 4°C in a swing arm rotor centrifuge. The cells were lysed in 50µl of cold ATAC Resuspension Buffer (10mM Tris-HCl pH 7.5, 10mM NaCl, 3mM MgCl_2_) containing 0.1% IGEPAL-630, 0.1% Tween-20, and 0.01% digitonin and incubated on ice for 3 minutes. The lysate was topped up with 1ml cold ATAC Resuspension Buffer containing 0.1% Tween-20 and underwent centrifugation at 500g for 10 min at 4°C in a swing arm rotor centrifuge. The pellet was resuspended in 50µl of transposition mixture (1X Illumina Tagment DNA Buffer and 100nM Illumina TDE1 Tagment DNA Enzyme transposase (Illumina; 20034197), 0.33X PBS, 0.1% Tween-20, 0.01% digitonin) and incubated at 37°C for 30 minutes with 1000 RPM mixing. After undergoing purification with the Zymo DNA Clean & Concentrator-5 kit (Zymo Research; D4003), samples were eluted in 21µl of elution buffer, and combined with 2.5µl of 25µM i5 primer, 2.5µl of 25µM i7 primer (**Supplementary Table 7**), and 25µl 2X NEBNext High-Fidelity 2X PCR Master Mix (NEB; M0541S). A PCR was performed with the following conditions: 72°C for 5 min, 98°C for 30 sec, 8 cycles of [98°C for 10 sec, 63°C for 30 sec, 72°C for 1 min]. The samples underwent a second Zymo DNA Clean & Concentrator-5 purification and were eluted in 30µl of elution buffer prior to a 1.2X AMPure XP beads (Beckman Coulter; A63881) cleanup. Following QC on a Bioanalyzer, libraries were multiplexed and sequenced (paired-end 50 bp) using a HiSeq 2000 instrument (Illumina).

### CUT&Tag

CUT&Tag was performed as described previously^35^ with minor modifications using the EpiCypher protocol. hiPSCs were washed once with DPBS (Life Technologies; 14190144) and incubated with Accutase (StemCell Technologies; 07922) for 5 min at 37°C. The cells were then centrifuged for 3 min at 300 x g at RT, resuspended in 1X DPBS and counted. 2 x 10^5^ cells per CUT&Tag reaction were centrifuged for 3 min at 300 x g at RT and then resuspended in 100 μl/sample of cold NE Buffer (20 mM HEPES–KOH, pH 7.9, 10 mM KCl, 0.1% Triton X-100, 20% Glycerol, 0.5 mM Spermidine (Sigma-Aldrich, 05292), 1x Roche cOmplete^TM^, Mini, EDTA-free Protease Inhibitor) and incubated for 10 min on ice. The nuclei were centrifuged for 3 min at 600 x g at RT, resuspended in cold NE Buffer to achieve a final concentration of 1×10^6^ nuclei/ml. 1 x 10^5^ nuclei per reaction were subjected to CUT&Tag.

The total volume of concanavalin A (ConA) beads (EpiCypher, 21-1401) of 11 μl per sample was transferred to a 1.5 ml tube for batch processing. ConA beads were washed twice on a magnetic stand with 100 μl/sample Bead Activation Buffer (20 mM HEPES, pH 7.9, 10 mM KCl, 1 mM CaCl_2_, 1 mM MnCl_2_) and resuspended in a final volume of 11 μl per sample. 10 μl beads were aliquoted into separate tubes for each sample and incubated with 100 μl of nuclei for 10 min at RT. The tubes were placed on a magnet separator (DynaMag-PCR; ThermoFisher Scientific), and the supernatant was discarded. Beads were then resuspended in 50 μl of cold Antibody150 Buffer (20 mM HEPES, pH 7.5, 150 mM NaCl, 0.5 mM Spermidine, 1x Roche cOmplete^TM^, Mini, EDTA-free Protease Inhibitor, 0.01% digitonin (Sigma-Aldrich, D141), 2mM EDTA) containing primary antibody in a 1:50 dilution (anti-H3K9me3, Active Motif 39062; anti-CTCF, Active Motif 61932; anti-H3K122ac, Abcam ab33309; anti-H4K12ac, Diagenode C15410331-10). Samples were incubated overnight at 4°C on an orbital shaker.

The following day, the tubes were placed on a magnet separator, and the supernatant was discarded. The beads were incubated in Digitonin150 Buffer (20 mM HEPES, pH 7.5, 150 mM NaCl, 0.5 mM Spermidine, 1x Roche cOmplete^TM^, Mini, EDTA-free Protease Inhibitor, 0.01% digitonin) containing 0.5 μg of secondary antibody (anti-rabbit Epicypher; 13-0047) for 1 hour at RT. The beads were then washed twice with Digitonin150 Buffer and then resuspended in 50 μl of Digitonin300 Buffer (20 mM HEPES, pH 7.5, 300 mM NaCl, 0.5 mM Spermidine, 1x Roche cOmplete^TM^, Mini, EDTA-free Protease Inhibitor, 0.01% digitonin), and 1.25 μl of CUTANA pAG-TN5 (EpiCypher, 15-1017) was added. After incubation at RT for 1 hour, the tubes were placed on a magnet and washed twice with Digitonin300 Buffer.

The beads were then resuspended in 50 μl of chilled Tagmentation Buffer (Digitonin300 Buffer, 10 mM MgCl_2_) and incubated for 1 hour at 37°C in thermocycler. The tube was placed on a magnetic separator and the supernatant was discarded. The beads were washed once with 50 μl RT TAPS Buffer (10 mM TAPS pH 8.5, 0.2 mM EDTA). The beads were resuspended in 5 μl of RT SDS Release Buffer (10 mM TAPS, pH 8.5, 0.1% SDS) and vortexed at maximum speed for 10 seconds followed by a brief centrifugation. The beads were then incubated for 1 hour at 58°C in a thermocycler. Following incubation, 15 μl of RT SDS Quench Buffer (0.67% Triton-X 100 in molecular grade H_2_O) was added to the beads and vortexed at maximum speed for 20 seconds, followed by a brief centrifugation.

For library amplification, 1 μl of each barcoded i5 and i7 primers (10μM stock) (**Supplementary Table 7**) was used and PCR was performed using CUTANA High Fidelity PCR mix (EpiCypher, 15-1018). The following PCR programme was used for all libraries: 58°C for 5 min, 72°C for 5 min, 98°C for 45 s, 11-13 cycles of 98°C for 15 s, 60°C for 10 s, 72°C for 1 min. Libraries were purified twice using 1X volume of AMPure XP Beads (Beckman Coulter; A63881) and resuspended in 15 μl of 0.1X TE buffer. Following QC on a Bioanalyzer, libraries were multiplexed and sequenced (paired-end 150 bp) using a NovaSeq 6000 instrument (Illumina).

### STARR-seq plasmid library preparation

Genomic DNA was purified from cultured cells (Monarch Genomic DNA Purification Kit, NEB, T3010S). 2 µg of purified DNA was sonicated targeting an 800 bp peak (E220 Focused-ultrasonicator, Covaris), and subjected to double-sided AMPure XP bead (Beckman Coulter, A63881) size selection between ratios 0.5× and 0.6× aiming at a fragment size range between 700 and 900 bp. Fragment sizes before and after selection were analysed on an agarose gel. Size selected DNA was end-repaired and ligated with the NEBNext hairpin adaptor (NEBNext Ultra II DNA Library Prep Kit for Illumina, NEB, E7645S, and NEBNext Multiplex Oligos for Illumina, NEB, E7335S). After adaptor ligation, the DNA was purified by 0.8× AMPure XP bead cleanup (Beckman Coulter, A63881) and eluted in 100 µL buffer EB (QIAGEN, 19086) in total. To generate the insert library for cloning, the adaptor ligated DNA was amplified with the STARR-seq library cloning forward (TAGAGCATGCACCGGACACTCTTTCCCTACACGACGCTCTTCCGATCT) and reverse (GGCCGAATTCGTCGAGTGACTGGAGTTCAGACGTGTGCTCTTCCGATCT) primers^32^, which added 15 bp homology sequence to both 3’ and 5’ ends. 20 PCRs (program: 45 s 98°C, 8 cycles of (15 s 98°C, 30 s 65°C, 45 s 72°C), 120 s 72°C) were performed in total for all the adaptor ligated DNA, where each 50 µL reaction contained 5 µL of the DNA, 2.5 µL library cloning forward primer (10 μM), 2.5 µL library cloning reverse primer (10 μM), 25 µL KAPA 2× HiFi HotStart Ready Mix (Roche, KK2602) and 15 µL nuclease-free water (Invitrogen, AM9938). The reactions were pooled, purified by 0.8× AMPure XP beads (Beckman Coulter, A63881) and eluted in 100 µL buffer EB (QIAGEN, 19086) in total. 40 µg hSTARR-seq_ORI vector (Addgene, 99296) was digested with 800 units of AgeI-HF (NEB, R3552L) and 800 units of SalI-HF (NEB, R3138L) in 16 of 50 µL reactions for 2 h at 37°C followed by deactivation for 20 min at 65°C. The reactions were size selected on an agarose gel for the approximately 3000 bp band, and the selected plasmids were eluted from the gel using QIAquick Gel Extraction kit (QIAGEN, 28704) into 400 µL buffer EB (QIAGEN, 19086) in total. The eluted plasmids were pooled, purified by 1.8× AMPure XP beads (Beckman Coulter, A63881) and eluted in 50 µL buffer EB (QIAGEN, 19086). To clone the insert library into the digested plasmid vector, 0.0667 pmol vector and 0.1333 pmol insert library were processed by each NEBuilder HiFi DNA Assembly (NEB, E2621L) reaction, where 40 reactions were performed in total. The reactions were pooled and purified by 1.8× AMPure XP beads (Beckman Coulter, A63881) and eluted in 30 µL nuclease-free water (Invitrogen, AM9938). The cloned plasmid concentration was measured by NanoDrop Spectrophotometer (Thermo Scientific). If the concentration was lower than 100 ng/µL, more cloning reactions were performed and the products were pooled and concentrated by 1.8× AMPure XP beads (Beckman Coulter, A63881) purification. To transform MegaX DH10B T1R Electrocomp Cells (Invitrogen, C640003) with the cloned plasmids, 20 electroporation reactions were performed on the Gene Pulser II Electroporation System (Bio-Rad). One day before electroporation, 12 L of LB media was pre-warmed at 37°C; 10 µL and 20 µL pipette tips, 1.5 mL DNA LoBind tubes (Eppendorf, 0030108051) and Gene Pulser/MicroPulser Electroporation Cuvettes, 0.1 cm gap (Bio-Rad, 1652083) were pre-cooled to 4°C. One hour before electroporation, at least 24 mL Recovery Media (provided in the MegaX DH10B T1R Electrocomp Cells kit, Invitrogen, C640003) was warmed up to 37°C. The following preparations were done in a 4°C cold room. At least 460 µL bacteria were thawed on ice, only bacteria with at most one freeze-thaw cycle were accepted for use. For each reaction, 100 ng plasmids in less than 1 µL nuclease-free water were added to 20 µL bacteria, mixed by gently flicking the bottom of the tube a few times, and carefully transferred to a Gene Pulser/MicroPulser Electroporation Cuvette, 0.1 cm gap (Bio-Rad, 1652083). The cuvette was tapped on the bench a few times to remove any air bubbles down the bottom of the cuvette, and left on ice until electroporation. One pUC19 control reaction using 1 µL pUC19 DNA (provided in the MegaX DH10B T1R Electrocomp Cells kit, Invitrogen, C640003) and 20 µL bacteria was prepared in addition to the 20 STARR-seq reactions. The cuvettes were then subjected to electroporation under the following settings on the Gene Pulser II Electroporation System (Bio-Rad), 2 kV, 25 µF, 200 Ω, 1 mm. Any result with a measured actual voltage of 2±0.1 kV and a measured time constant of 4.9±0.2 msec was regarded as a successful electroporation. After each electroporation reaction, 1 mL of pre-warmed Recovery Medium (provided in the MegaX DH10B T1R Electrocomp Cells kit, Invitrogen, C640003) was immediately added to the cuvette. Electroporated bacteria were resuspended into the medium and transferred to a 14 mL polypropylene round-bottom tube (Falcon, 352059). The bacteria were recovered for 1 h at 37°C with constant shaking at 200 rpm. All STARR-seq transformed bacteria were pooled. To estimate the transformation efficiency, 100 µL of 1:50, 1:500, and 1:5000 diluted STARR-seq transformed bacteria, and 1:50, 1:500 diluted pUC19 transformed bacteria were plated on selective LB agar plates containing 100 μg/mL ampicillin. After 16 h incubation at 37°C, more than 50 colonies found on both the 1:5000 diluted STARR-seq plate and the 1:500 diluted pUC19 plate were signs for successful transformation. The rest of the pooled STARR-seq transformed bacteria were evenly added to 12 L of pre-warmed LB with 100 μg/mL ampicillin, and incubated for 12 to 14 h at 37°C with constant shaking at 200 rpm. The bacteria culture was harvested by centrifugation and the pellets were stored at −20°C. The pellets were purified using the QIAGEN Plasmid Plus Giga Kit (QIAGEN, 12991), eluted in 3 mL buffer EB (QIAGEN, 19086), further purified and concentrated by 1.8× AMPure XP beads (Beckman Coulter, A63881) and eluted in nuclease-free water (Invitrogen, AM9938) at a final concentration of 5 μg/μL. To generate sequencing library for the STARR-seq plasmid library, 10 PCRs (program: 45 s 98°C, 9 cycles of (15 s 98°C, 30 s 65°C, 45 s 72°C), 120 s 72°C) were performed, where each 50 μL reaction contained 100 ng plasmid library, 2.5 μL Illumina i5 primer (10 μM), 2.5 μL Illumina i7 primer (10 μM), 25 µL KAPA 2× HiFi HotStart Ready Mix (Roche, KK2602) and nuclease-free water (Invitrogen, AM9938). The amplified sequencing libraries were pooled and purified by 0.8× AMPure XP beads (Beckman Coulter, A63881) and eluted in 20 µL buffer EB (QIAGEN, 19086). Library concentration and quality were analysed using the Agilent High Sensitivity DNA Kit (Agilent, 5067-4626) on a 2100 Bioanalyzer (Agilent).

### STARR-seq enhancer annotation

At least 250 million hiPSCs (HPSI0215i-yoch_6) were prepared for each biological replicate. Cells were transfected in the MaxCyte R-1000 Processing Assemblies (MaxCyte, ER001M1-10) for the STARR-seq samples and in the R-50×3 Processing Assemblies (MaxCyte, ER050U3-10) for the control samples, both on the MaxCyte ATx electroporation platform (MaxCyte). Cells were harvested, washed by Dulbecco’s Phosphate Buffered Saline (DPBS, Cytiva HyClone, SH30028.02) and counted for cell number and viability. 200 million cells were taken for the STARR-seq sample, and 10 million cells were taken for each of the three control samples: non-electroporated negative control, electroporated without DNA negative control, and GFP-transfected positive control (GFP-expressing plasmid; MaxCyte). Cells were centrifuged for 3 min at 300 g, and the supernatant was removed. Cells of the non-electroporated negative control were directly plated in culture. 40 μL of STARR-seq plasmid library at a concentration of 5 μg/μL was added to the STARR-seq sample cell pellet; 2 μL of GFP plasmid at a concentration of 5 μg/μL was added to the positive control cell pellet. Cells with or without plasmid DNA were then resuspended in MaxCyte Electroporation Buffer (provided in the Processing Assembly kits) to a final volume of 1 mL for the STARR-seq sample and 50 μL for the controls. The samples were then transferred to the corresponding Processing Assemblies (MaxCyte, ER001M1-10 and MaxCyte, ER050U3-10) and subjected to electroporation under the “optimisation 7” program of the ATx electroporation platform controlling software (MaxCyte). After electroporation, cells were transferred to uncoated culture vessels with excess surface area, aiming for maximum exposure to air, and recovered for 30 min at 37°C and 5% CO_2_. Recovered cells were then diluted to 1 million cells per mL of media with supplement and 10 µM Rho-associated protein Kinase (ROCK) inhibitor (Cell Guidance Systems, Y-27632), and cultured on vitronectin-coated (Thermo Fisher Scientific, A14700) vessels. For the STARR-seq sample, cells were cultured for 6 h before being harvested for total RNA purification. For the control samples, cells were cultured for 24 h, then in fresh media with supplement without ROCK inhibitor for another 24 h, before being harvested for flow cytometry analysis on GFP expression to determine the transfection efficiency of the experiment. For the STARR-seq sample, total RNA was purified from the harvested cells using the QIAGEN RNeasy Maxi Kit (QIAGEN, 75162), and eluted in three consecutive 1 mL elutions in RNase-free water (provided in the RNeasy Maxi Kit). An aliquot of the total RNA was taken for integrity analysis using the Agilent RNA 6000 Pico Kit (Agilent, 5067-1513) on a 2100 Bioanalyzer (Agilent). The rest of the total RNA was stored at −80°C until sequencing library preparation. Any elution fractions with concentrations greater than 100 ng/μL were used for sequencing library preparation. mRNA was isolated from the total RNA using Dynabeads™ Oligo(dT)_25_ (Invitrogen, 61002) and eluted in 200 μL buffer EB (QIAGEN, 19086). The isolated mRNA concentration was measured by NanoDrop Spectrophotometer (Thermo Scientific). The mRNA was subjected to DNA digestion by TURBO DNase (Invitrogen, AM2238). Each 100 μL reaction contained at most 20 μg mRNA, 2.4 μL TURBO DNase and 1× TURBO DNase buffer (provided with the TURBO DNase). The reactions were pooled, purified by 1.8× RNAClean XP beads (Beckman Coulter, A63987) and eluted in 200 μL nuclease-free water (Invitrogen, AM9938) in total. cDNA was synthesised from the mRNA by SuperScript IV Reverse Transcriptase (Invitrogen, 18090200), supplemented by Murine RNase Inhibitor (40,000 U/mL, NEB, M0314L). The number of reactions was determined by the mRNA yield, according to 500 ng mRNA per 20 μL reaction. The reverse transcriptase reactions were followed by RNaseA treatment, where 0.2 μL RNaseA (10 mg/mL, Thermo Fisher, EN0531) was added to each reverse transcriptase reaction and the mixture was incubated for 1 h at 37°C. The reactions were purified by 1.8× AMPure XP beads (Beckman Coulter, A63881), eluted in 2 µL buffer EB (QIAGEN, 19086) per reverse transcriptase reaction and pooled. To enrich STARR-seq-specific cDNA, the purified cDNA was subjected to junction PCR, where one reaction amplified up to 20 µL purified cDNA. Each 50 μL reaction (program: 45 s 98°C, 16 cycles of (15 s 98°C, 30 s 65°C, 60 s 72°C), 120 s 72°C) contained the cDNA, 2.5 μL junction forward primer (10 μM, TCGTGAGGCACTGGGCAG*G*T*G*T*C), 2.5 μL junction reverse primer (10 μM, CTTATCATGTCTGCTCGA*A*G*C)^32^, 25 µL KAPA 2× HiFi HotStart Ready Mix (Roche, KK2602) and buffer EB (QIAGEN, 19086). The junction PCRs were purified by 0.8× AMPure XP beads (Beckman Coulter, A63881), eluted in 20 µL buffer EB (QIAGEN, 19086) per reaction and pooled. To generate the sequencing library, the junction PCR product was further subjected to sequencing-ready PCR. Each biological replicate used a unique set of 16 Illumina i5 and i7 dual-indexing combinations, and the junction PCR product was evenly split into 16 according to one reaction per dual-indexing combination. Each 50 μL reaction (program: 45 s 98°C, 8 cycles of (15 s 98°C, 30 s 65°C, 45 s 72°C), 120 s 72°C) contained the purified junction PCR product, the corresponding 2.5 μL Illumina i5 primer (10 μM) and 2.5 μL Illumina i7 primer (10 μM), 25 µL KAPA 2× HiFi HotStart Ready Mix (Roche, KK2602) and buffer EB (QIAGEN, 19086). The reactions were pooled, purified by 0.8× AMPure XP beads (Beckman Coulter, A63881) and eluted in 40 µL buffer EB (QIAGEN, 19086) in total. Library concentration and quality were analysed using the Agilent High Sensitivity DNA Kit (Agilent, 5067-4626) on a 2100 Bioanalyzer (Agilent).

### Promoter Capture Hi-C

PCHi-C was performed as described^37^ in the HipSci lines HPSI1113i-bima_1, HPSI0114i-kolf_2, HPSI0214i-kucg_2, HPSI1113i-podx_1, HPSI1113i-qolg_1 (all DpnII), and HPSI0114i-kolf_2, HPSI1113i-podx_1, HPSI1113i-qolg_1 (all HindIII) with the following modifications: after Klenow-mediated fill-in of restriction fragments overhang (with biotin-dATP, dCTP, dGTP and dTTP), nuclei were spun down for 5 minutes (2500 rpm at 4°C). The supernatant was removed, and the nuclei pellet was resuspended in 950 µl 1x ligation buffer (supplied with T4 DNA ligase Invitrogen 15224-025), before 50 µl of T4 DNA ligase (Invitrogen 15224-025) was added for the ligation for 4 hours at 16°C.

### Luciferase assay plasmid preparation

20-base homology sequences were added to both ends of the selected candidate sequences, positive and negative enhancer controls for cloning purposes (upstream: CTGGCCTAACTGGCCGGTAC, downstream: CCTCGAGGATATCAAGATCT). The combined sequences were artificially synthesised as gBlocks Gene Fragments (Integrated DNA Technologies). Minimal Promoter-Driven Firefly Luciferase Vector (pGL4.23[luc2/minP] Vector, Promega, E8411) and Promoterless Renilla Luciferase Basic Vectors (pGL4.70[hRluc] Vector, Promega, E6881) were amplified by transformation into NEB 5-alpha Competent E. coli (High Efficiency) (NEB, C2987H) and minipreps using QIAprep Spin Miniprep Kit (QIAGEN, 27104). For each candidate and positive and negative enhancer controls, 500 ng empty firefly luciferase vector (pGL4.23) was restricted by KpnI-HF (NEB, R3142S) and NheI-HF (NEB, R3131S), purified by 0.8× AMPure XP beads (Beckman Coulter, A63881) and eluted in buffer EB (QIAGEN, 19086). To clone the gBlocks fragment insert into digested plasmid vector, for each candidate, 0.0667 pmol vector and 0.1333 pmol insert were processed by NEBuilder HiFi DNA Assembly (NEB, E2621L) reaction. After the reaction was completed, 1 out of the 20 µL reaction mixture was used to transform the cloned plasmid into 50 µL NEB 5-alpha Competent E. coli (High Efficiency) (NEB, C2987H) by heat shock at 42°C for 30 s. After 5 min incubation on ice, the transformed bacteria were diluted in 1 mL LB medium in a 14 mL polypropylene round-bottom tube (Falcon, 352059) and recovered for 1 h at 37°C with constant shaking at 200 rpm. 1 µL of the recovery mixture was then diluted in 100 µL LB media and plated on a selective LB agar plate containing 100 μg/mL ampicillin. After 16 h incubation at 37°C, a single colony was inoculated into 2 mL LB media containing 100 μg/mL ampicillin and incubated in a 14 mL polypropylene round-bottom tube (Falcon, 352059) for 12 to 14 h at 37°C with constant shaking at 200 rpm. The amplified cloned plasmid was purified from the bacteria using the QIAprep Spin Miniprep Kit (QIAGEN, 27104) and eluted in 50 μL buffer EB (QIAGEN, 19086). The plasmid was further purified and concentrated by 1.8× AMPure XP beads (Beckman Coulter, A63881) and eluted in nuclease-free water (Invitrogen, AM9938) at a final concentration of 1 μg/μL. To check the plasmid integrity, 100 ng of the empty firefly, Renilla and all cloned candidate and control plasmids were restricted by ClaI (NEB, R0197S) and analysed on an agarose gel.

### Luciferase assay

The minimum number of cells required for each biological replicate experiment was calculated as follows, 1 million hiPSC per sample * total number of samples (the number of candidates + 4 controls). The four controls were no DNA control, firefly luciferase empty plasmid control, positive enhancer control and negative enhancer control. Cells were harvested, washed by Dulbecco’s Phosphate Buffered Saline (DPBS, Cytiva HyClone, SH30028.02) and counted for cell number and viability. 1 million cells were transferred to an individual 14 mL polypropylene round-bottom tube (Falcon, 352059) for each of the samples and controls. Cells were centrifuged for 3 min at 300 g, and the supernatant was removed. 1 μL of 1 μg/μL candidate/control plasmid and 1 μL of 1 μg/μL Renilla luciferase plasmid were added to the corresponding cell pellet except the no DNA control. To transfect cells using the 4D-Nucleofector (Lonza) core and X units and the P3 Primary Cell 4D-Nucleofector X Kit L (Lonza, V4XP-3024), for each sample cells and plasmid DNA were mixed (cells only for the no DNA control) and resuspended in 100 μL Nucleofection Solution (provided in the P3 kit, Lonza, V4XP-3024) and transferred to a Single Nucleocuvette (100 µl, provided in the P3 kit, Lonza, V4XP-3024). The cells were then subjected to 4D-Nucleofector X Unit (Lonza) and electroporated using the CA137 program. After nucleofection the cells were recovered within the Nucleocuvette at room temperature for 10 min, diluted in 1 mL media with supplement and 10 µM Rho-associated protein Kinase (ROCK) inhibitor (Cell Guidance Systems, Y-27632) and cultured on one well of a vitronectin- (Thermo Fisher Scientific, A14700) coated 12-well plate for 24 h. The media was then removed and cells were washed on-plate using Dulbecco’s Phosphate Buffered Saline (DPBS, Cytiva HyClone, SH30028.02) and lysed on-plate by adding 250 μL 1× Passive lysis Buffer (provided in Dual-Luciferase® Reporter Assay System, Promega, E1910) and incubating at room temperature for 15 min under constant rocking. The firefly and Renilla luciferase expression levels were measured by luminescence using the Dual-Luciferase® Reporter Assay System (Promega, E1910) in wells of 96-well plates (96-well EIA/RIA Clear Flat Bottom Polystyrene High Bind Microplate, Individually Wrapped, without Lid, Nonsterile, Corning, 3590) in a FLUOstar Omega microplate reader (BMG Labtech).

### RNA-seq

1 to 5 million cells were prepared for each biological replicate. Cells were harvested and washed by Dulbecco’s Phosphate Buffered Saline (DPBS, Cytiva HyClone, SH30028.02). Total RNA was purified from the cell pellet using the Monarch Total RNA Miniprep Kit (NEB, T2010S). An aliquot of the total RNA was taken for integrity analysis using the Agilent RNA 6000 Pico Kit (Agilent, 5067-1513) on a 2100 Bioanalyzer (Agilent). The rest of the total RNA was stored at −80°C until sequencing library preparation. RNA-seq libraries were prepared using the NEBNext Ultra II Directional RNA Library Prep Kit for Illumina (NEB, E7760L), NEBNext Poly(A) mRNA Magnetic Isolation Module (NEB, E7490L) and NEBNext Multiplex Oligos for Illumina (Index Primers Set 1, 2, 3 and 4, NEB, E7335S, E7500S, E7710S, E7730S), where for each preparation 500 ng total RNA was used as the starting material and 8 cycles were used for the final sequencing-ready PCR. Library concentration and quality were analysed using the Agilent High Sensitivity DNA Kit (Agilent, 5067-4626) on a 2100 Bioanalyzer (Agilent). The STARR-seq sequencing libraries were sequenced in NextSeq500 HighOutput 75 bp Paired End runs and HiSeq2500-RapidRun 50bp Paired End runs on corresponding sequencing platforms (Illumina). The RNA-seq sequencing libraries were sequenced in HiSeq2500-RapidRun 50bp Paired End runs on the HiSeq sequencing platform (Illumina).

### Transient transcriptome sequencing (TT-seq)

TT-seq was performed as previously described^79^ with minor modifications. hiPSCs were labelled with 500 μM 4-thiouridine (Sigma) for 20 min. Cells were counted and collected in TRIzol. RNA was chloroform-extracted, DNase-treated, and chloroform-extracted. To 60 μg total RNA was added 2x fragmentation buffer (final concentration: 75 mM Tris-Cl, pH 8.3, 112.5 mM KCl, and 4.5 mM MgCl_2_) and heated to 95 °C for 5 min. Fragmentation was stopped by adding EDTA to 50 mM and placing samples on ice. RNA was ethanol-precipitated and resuspended in H_2_O. Fragment sizes were checked on the Bioanalyser (peak size of ∼800 nucleotides). The biotinylation reaction was performed with 0.025 mg/mL MTSEA-Biotin XX (Biotum) in reaction buffer (20% N,N-dimethyl-formamide, 1 mM EDTA, 10 mM HEPES pH 8) for 45 min in the dark. Labelled RNA was chloroform-extracted ethanol-precipitated and resuspended in 90 μL H_2_O. 75 microliters of DynabeadsTM M-280 streptavidin (Thermo Fisher) were prewashed with decon solution (0.1 M NaOH + 50 mM NaCl), 2x 100 mM NaCl, 2x high salt buffer (100 mM Tris-Cl pH 7.4, 10 mM EDTA pH 8, 1 M NaCl, 0.05% (vol/vol) Tween 20), and resuspended in high salt buffer. Labelled RNA was denatured by heating it to 65 °C for 5 min and placing it on ice for 2 min. Ten microliters of high salt buffer were added to labelled RNA. Prewashed beads were placed on the magnet, the supernatant discarded, and then beads were resuspended in the labelled RNA. Beads and RNA were rotated for 30 min in the dark. Beads were then washed 4x for 1 min with high salt buffer. Labelled RNA was eluted 2x with 100 mM DTT (1,4-dithiothreitol). RNA was cleaned with a RNeasy MinElute Clean-up Kit (Qiagen Cat. No 74204). 250 ng of RNA was used for library preparation using the CORALL total RNAseq library preparation kit (Lexogen Cat. no. 147.24) using eight cycles of PCR amplification. Libraries were sequenced in Novaseq 6000 (Novogene) with 150 bp paired-end reads.

### Immunofluorescence

HiPSC colonies were cultured on 16 mm glass coverslips (Fisherbrand™, 12323148) placed in 12-well plates. After the induction period, the colonies were washed with PBS and then fixed in 4% PFA for 20 minutes at room temperature (RT). Following three washes with 1X PBS, the colonies were incubated in a permeabilization and blocking buffer (PB; 1% BSA, 0.1% Triton X-100 in PBS) for 30 minutes at RT. The cells were incubated overnight at 4°C with primary antibodies against SOX2 (Santa Cruz, sc-365823, 1:500) and NANOG (R&D -AF1997, 1:100) in PB. The following day, they were incubated for one hour at RT with fluorescence-conjugated secondary antibodies (Thermo Fisher Scientific, A32787, A11057, 1:400), followed by three washes with 1X PBS. Finally, the colonies were mounted with ProLong™ Glass Antifade Mountant with NucBlue™ Stain (Thermo Fisher Scientific, P36983) and allowed to dry. The samples were imaged using a Leica Stellaris 8 confocal microscope with LAS X software and analyzed with CellProfiler 4.2.6 and ImageJ 1.54f software.

### Live cell imaging

Live cell imaging was performed using the Sartorius Incucyte S5 system, and time-lapse images and videos were analysed using the Sartorius Incucyte 2022B Rev2 software interface.

### RT–qPCR

Total RNA was extracted using the Monarch® Total RNA Miniprep Kit (NEB, T2010S), and 1 µg of RNA was used for complementary DNA synthesis with the RevertAid First Strand cDNA Synthesis Kit (Thermo Fisher Scientific, K1622) according to the manufacturer’s protocols. qPCR was performed on a CFX Opus 96 Real-Time PCR System (Bio-Rad) using specific primers for target genes (**Supplementary Table 7**) and qRT-PCR Brilliant III SYBR Master Mix (Agilent, 600886). The qPCR cycle conditions were as follows: 40 cycles of 10 seconds at 95°C, 30 seconds at 55°C, and 30 seconds at 72°C. Primers were designed using NCBI Primer3 and synthesised by Integrated DNA Technologies. Quantifications were calculated and normalized to the endogenous GAPDH internal control. The ΔΔCt method was used for relative quantification of each target gene compared to an internal control.

### CRISPRa

We designed single guide RNA (sgRNA) and used CRISPRa plasmids based on the doxycycline (DOX)-inducible catalytically inactive Cas9 (dCas9) as described previously^16^. Three sgRNAs targeting the enhancer regions were designed using the gRNA design tool of Integrated DNA Technologies (**Supplementary Table 8**). 1.5 ug of each of the piggyBac-based CRISPRa (puromycin-resistant cassette; Addgene, 183409), sgRNA plasmids (neomycin-resistant cassette; Addgene, 183411), and a hyperactive piggyBac transposase plasmid were co-transfected into hiPSC lines using the Invitrogen Neon Transfection System. Increasing concentration of Puromycin (up to 1 μg/ml) and Neomycin (up to 200 μg/ml) was used for selection of transgenic cells for 7–10 days. To induce the CRISPRa system and activation of the enhancers, DOX was used in 0.5 μg/ml concentration in TeSR-E8 medium for 4 days and cells were collected for further analysis after the induction period. Non-induced cells (without DOX) were used as a negative control.

### CRISPRi

CRISPRi was carried out as previously described^16^. 1.5 μg of piggyBac-based CRISPRi plasmid (Addgene, 183410), three sgRNA plasmids for each region (**Supplementary Table 8**), and piggyBac-transposase plasmid were co-transfected into hiPSCs using the Invitrogen Neon Transfection System. Stable cell lines were then selected with increasing concentrations of puromycin and neomycin for 7– 10 days after transfection. To test the effect of enhancer perturbation on the expression of target gene(s), CRISPRi hiPSC lines were induced with 0.5 μg/ml of DOX and 10 μM of Trimethoprim (TMP) to induce and stabilise dCas9-KRAB for 4 days and collected for further analysis. Non-induced cells (without DOX and TMP) were used as a negative control.

### RNA-seq data processing and analyses

RNA-seq data analysis and quality check of the raw reads were done by FastQC v0.11.8 (www.bioinformatics.babraham.ac.uk/projects/fastqc) and subsequently reads were aligned to the human reference genome GRCh38 using STAR v2.5.2b2^80^. Duplicates were removed using Picard (v2.9.0) (https://broadinstitute.github.io/picard/). Aligned reads were assigned to gene annotation using HTSeq-count version 0.10^81^. Differential analysis and normalization were performed using DESeq2^82^. Genes were considered differentially expressed if they had an absolute log2fold change exceeding 1 and a *Padj* of less than 0.01. PCA was performed by prcomp function and plotted by ggplot2 in R^83^. Heatmaps were generated using the pheatmap R package (https://rdrr.io/cran/pheatmap/). GO enrichment analysis was performed using clusterProfiler^84^.

### TT-seq data processing and analyses

TT-seq data analysis and quality check of the raw reads were done by FastQC v0.11.8 (www.bioinformatics.babraham.ac.uk/projects/fastqc) and subsequently reads were aligned to the human reference genome GRCh38 using STAR v2.5.2b2^80^. The BAM files generated for biological replicates were merged with samtools, merge and filtering for only uniquely mapped reads using the samtools flag -q 255 followed by sorting and indexing. Coverage tracks requiring bigWig files and heatmaps were generated using deepTools v3.3.1^85^.

### ATAC-seq data processing and analyses

ATAC-seq data analysis and quality inspection was performed using FastQC v0.11.5 (www.bioinformatics.babraham.ac.uk/projects/fastqc). Nextera Transposase adapter and low-quality bases were eliminated using Cutadapt v1.17^86^. After trimming, reads were mapped to human genome GRCh38 assembly using bowtie2 v2.3.4.2^87^. Mapped pairs were further filtered to maintain mapping quality above 20, using custom scripts and SAMtools v1.9^88^. PCR duplicates and mitochondrial reads were removed by Picard (https://broadinstitute.github.io/picard/) and samtools as well as reads blacklisted by The Encyclopedia of DNA Elements (ENCODE) project^89^. Peaks of accessible chromatin were identified for each sample using MACS2 v2.1.2^90^ with the following settings: -f BEDPE --keep-dup (51). Coverage tracks requiring bigWig files and heatmaps were generated using deepTools v3.3.1^85^.

### CUT&Tag data processing and analyses

CUT&Tag datasets were processed and mapped to the human reference genome GRCh38 as described previously^35^. After mapped reads were further filtered to maintain mapping quality above 30, using custom scripts and SAMtools v1.9^88^, peaks were called using MACS2 v2.1.2^90^.. H3K9me3 peaks were called using the following settings: -broad -f BAMPE --keepdup all -p 1e-2. BigWig files required for coverage tracks and heatmaps were generated using deepTools v3.3.1^85^.

### ChIP-seq data processing and analyses

ChIP-seq datasets were mapped to the human reference genome GRCh38 using bowtie2 v2.3.4.2^87^. After mapped reads were further filtered to maintain mapping quality above 30, using custom scripts and SAMtools v1.9^88^, peaks were called using MACS2 v2.1.2^90^ with the following settings: narrowPeak representation was used for histone marks H3K4me3 and H3K27ac with the following settings: -f BAMPE --keepdup all -p 1e-2. H3K27me3 and H3K9me3 peaks were called using the following settings: -broad -f BAMPE --keepdup all -p 1e-2. BigWig files required for coverage tracks and heatmaps were generated using deepTools v3.3.1^85^. In order to separate enhancers into categories, we used a master set of peaks combined from several published ChIP-seq H3K27ac datasets (ME, TBL, MSC, NPC) from SRP000941. We called the peaks separately on the merged replicates in the respective studies. BigWig files required for coverage tracks and heatmaps were generated using deepTools v3.3.1^85^.

### STARR-seq data processing and analyses

Reads of input and output STARR-seq libraries were aligned to human genome GRCh38 by bowtie2^87^. Non-unique alignments and alignments with mapping quality less than 30 were filtered out. Duplicates were removed using Picard (v2.9.0) (https://broadinstitute.github.io/picard/).

For each output replicate, uniquely mapped distinct fragments from the 16 different indexing libraries were pooled and included in the downstream analysis. Duplicated fragments from different indexing libraries were treated as biological duplicates. We pooled five biological output replicates STARR-seq biological replicates were merged using samtools merge. STARR-seq peaks were called using STARRpeaker using default settings. BigWig files required for coverage tracks and heatmaps were generated using deepTools v3.3.1^85^.

### Promoter Capture Hi-C data processing and analyses

DpnII and HindIII PCHi-C data were mapped and filtered using HiCUP^50^ with the GCRh38 human genome build. BAM files were converted to a CHiCAGO-compatible format (.chinput) using bam2chicago script. CHiCAGO^51^ was used to define significant promoter interactions at the level of individual fragments. CHiCAGO uses a convolution background model for the background level of interactions for pooled baited and other regions and a weighted distance-dependent multiple testing correction.

Two biological replicates for each of 5 different cell lines were normalized and combined as part of the CHiCAGO pipeline. CHiCAGO interaction scores correspond to –log-transformed, weighted P-values for each fragment read pair. A CHiCAGO interaction score of 5 or above was considered significant based on previous empirical observations^51^. We used default CHiCAGO weights for HindIII data processing. To process DpnII data we applied fitdistcurve function from the CHiCAGO package to measure optimal weight parameters. PCHi-C contacts were visualized using WashU Browser (https://epigenomegateway.wustl.edu/).

### Identification of inter-allelic enhancer variants

SNP calling for allelic imbalance in STARR-seq data was performed following the GATK pipeline as recommended by the GATK Best Practices (software.broadinstitute.org/gatk/best-practices/) To begin with, all raw FASTQ files for each of the STARR-seq replicate was first converted to BAM format using the FastqToSam program. The bam files were combined using samtools merge and sorted by query name. MarkIlluminaAdapters was used to mark PCR duplicates and marked bam files were converted to interleaved FASTQ format using the SamToFastq program. FASTQ to Bam conversion was then performed using the BWA-MEM software and recalibrated after removing PCR duplicates. Haplotype calling was performed on each of the recalibrated BAM files using the HaplotypeCaller in discovery mode and combined using the CombineGVCFs program. Next, recalibration of SNPs and Indels was carried out using the VariantRecalibrator function using the HapMap v3.3 (priority = 15), 1000G_omni v2.5 (priority = 12), Broad Institute 1000G high confidence SNP list phase 1 (priority = 10), Mills 1000G golden standard INDEL list (priority = 12), and dbSNP v150 (priority = 2) databases. Post identification of SNPs that show allelic imbalance, N masked genome was generated using the SNPsplit_genome_preparation program. FASTQ files were then aligned to the N-masked genome and SNPsplit was used to separate reads into Allele1 and Allele2. Samtools pileup was further used to count read pileup at heterozygous SNP sites. Binomial testing was used to identify regions which show a significant (padj<0.05) difference in pileups between Allele1 and Allele2.

### Motif Discovery and Enrichment Analysis

The JASPAR 2022 core nonredundant motifs were used as the initial motif candidates list. Only transcription factor motifs which had at least 10 TPM levels of expression were selected for motif discovery analysis. AME^91^ from the MEME Suite (v5.5.1)^92^ was used to perform motif discovery and enrichment analysis. AME identifies known motifs that are relatively enriched in given sequences compared with control sequences.

### Published ChIP-seq data

All published data used in the analysis are available to download using following accession numbers: GSE69647 for naïve and primed hPSC H3K27ac, GSE59434 for primed hPSC H3K27me3 & H3K4me3, GSE75868 for primed hPSC H3K9me3, GSE19465 for primed hPSC H3K4me1, GSE51334 for primed hESC P300, GSE51334 for H1 hESC POLR2A, GSE16256 for primed hESC, MSC, NPC, TBL, ME.

### DeepSTARR

DeepSTARR is a deep learning model that directly predicts the activity of enhancers in DNA sequences^48^. To train and apply the model, we performed the pipeline as described previously^48^, with minor changes. We only included bins with more than five reads in the input and at least one read in the STARR-seq output. The genome was binned into regions of 499 bp with a stride of 100 bp. The included bins came from:

1. Random regions of accessible chromatin with low STARR-seq signal.
2. Random regions of accessible chromatin with high STARR-seq signal.
3. Random regions of inaccessible chromatin with various STARR-seq signals.

We augmented our dataset by adding the reverse complement of each original sequence with the same output. The model and performance evaluation were conducted as previously described^48^, with the exception of changing the input DNA sequence length to 499 bp.

## AUTHOR CONTRIBUTIONS

U.G. and S.S. conceived the project, planned and supervised experiments, interpreted data and wrote the manuscript, with input from all authors. Y.C. carried out STARR-seq, RNA-seq and luciferase enhancer reporter experiments. O.D. led all bioinformatic analyses, except for inter-allelic enhancer strength determination which was done by S.K. A.E. and L.W. carried out CRISPR experiments, supported by I.R.P. and J.K. A.E. performed RNA-seq, RT-qPCR and live and fixed cell imaging. F.T. and A.E. performed CUT&Tag experiments. S.B. carried out ATAC-seq and, together with S.S., PCHi-C. M.M.P. and M.P. performed TT-seq experiments and analysis.

## ACKNOWLEDGEMENTS

We are indebted to Marianna Romero for expert advice on MaxCyte electroporation protocols, to Simon Walker for expert help with fixed and live-cell imaging, and to Paula Kokko-Gonzales for optimising protocols for STARR-seq library sequencing. We thank Dominik Muehlen for advice on CUT&Tag and help with figure design, Kaia Spence for help with CRISPRi, Peter Rugg-Gunn for advice on hiPSC cell culture, Bernardo P. de Almeida for advice on DeepSTARR, and Olivia Cracknell for critical reading of the manuscript. S.S. was supported by a UKRI MRC Rutherford Fund Fellowship (MR/T016787/1) and a Career Progression Fellowship from the Babraham Institute. S.S., S.B. Y.C., and A.E. were supported by the Babraham Institute’s Strategic Programme Grant Epigenetics (BBS/E/B/000C0421). U.G. was supported by a Sofja Kovalevskaja Award of the Humboldt Foundation. O.D. and J.K. were enrolled in the Göttingen Graduate Center for Neurosciences, Biophysics, and Molecular Biosciences, supported with funds from the Sofja Kovalevskaja Award to U.G. and the International Max Planck Research School (IMPRS) for Genome Science (O.D.) and the IMPRS for Molecular Biology (J.K.). I.R.P. was supported by a Walter-Benjamin postdoctoral fellowship (DFG, 509071720) and a Klaus Tschira Boost Fund.

## DATA AVAILABILITY

Next generation sequencing data are available using GEO accession number XXXX.

---

**Figure S1:**
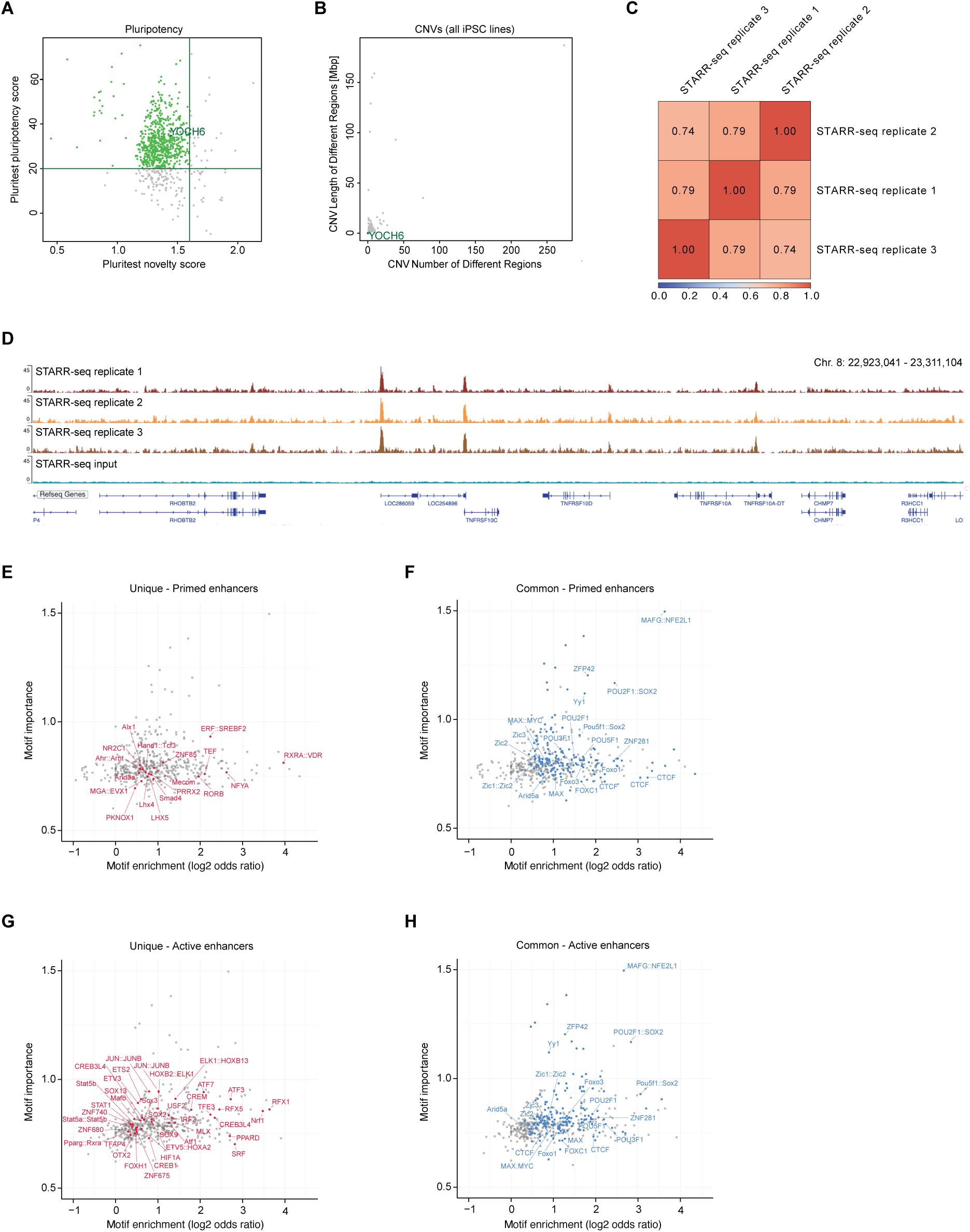
STARR-seq quality control and sequence motifs in active and primed enhancers. (A) Scatterplot showing Pluritest scores of HipSci hiPSC lines for pluripotency and novelty. The hiPSC line used for STARR-seq is highlighted (Yoch6; HPSI0215i-yoch_6). (B) Copy Number Variations (CNVs) in HipSci hiPSC lines. HiPSC line Yoch6 is highlighted. (C) Pearson correlations of STARR-seq replicates. (D) Example coverage tracks of STARR-seq replicates 1-3 and the STARR-seq input background control. (E-F) Scatter plot showing motif enrichment (x-axis) and DeepSTARR-predicted global importance (y-axis) at primed enhancers (red). Unique (E): Motifs called only at primed enhancers in comparison to active enhancers. Common (F): Motifs called at both enhancer classes. (G-H) Scatter plot showing motif enrichment (x-axis) and DeepSTARR-predicted global importance (y-axis) at active enhancers (red). Unique (G): Motifs called only at active enhancers in comparison to primed enhancers. Common (H): Motifs called at both enhancer classes.

**Figure S2:**
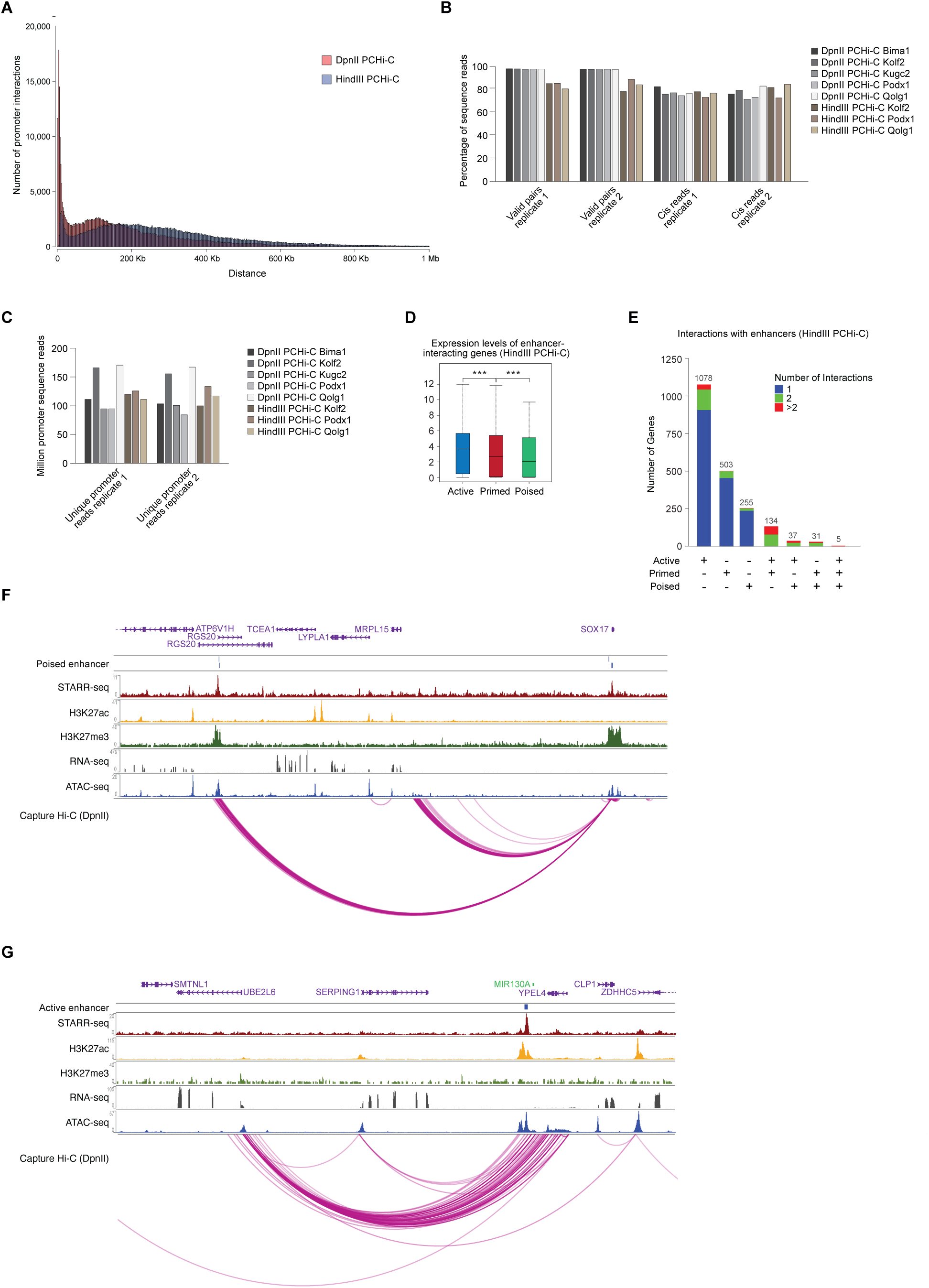
Promoter Capture Hi-C identifies target genes of enhancers. (A) Number of interactions and their distances to captured promoters in bp, in PCHi-C data generated using six- or four-base cutter restriction enzymes HindIII (HindIII PCHi-C) or DpnII (DpnII PCHi-C) with hiPSCs. (B) Percentage of valid and *cis* sequencing reads in HindIII or DpnII PCHi-C datasets with indicated hiPSC cell lines. (C) Number of promoter-centered reads (Unique promoter reads) in indicated PCHi-C datasets. (D) Expression (RNA-seq, log2 (normalised counts+1)) of enhancer-interacting genes. Interactions were mapped with HindIII PCHi-C data in hiPSCs. (E) Number of genes interacting with one, two or more than two active, primed and/or poised enhancers. Interactions were mapped using HindIII PCHi-C data. (F) Coverage tracks of genome-wide data on chromosome 8 showing a representative example of a poised enhancer. Capture Hi-C corresponds to DpnII PCHi-C data. (G) Coverage tracks of genome-wide data on chromosome 11 showing a representative example of an active enhancer. Capture Hi-C corresponds to DpnII PCHiC data.

**Figure S3:**
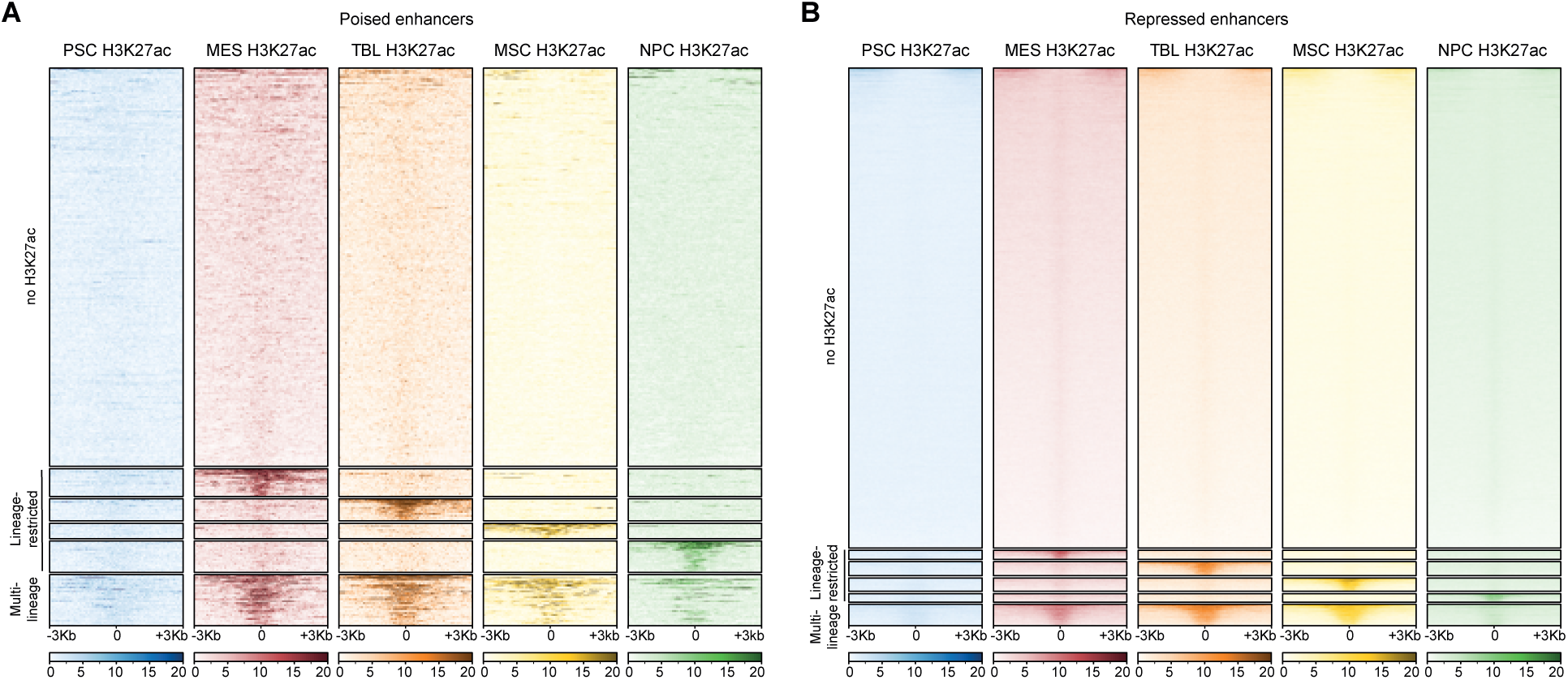
The majority of poised and chromatin-repressed enhancers in hiPSCs do not gain H3K27ac during differentiation. (A) Heatmap showing H3K27ac enrichment (ChIP-seq) at poised enhancers (mapped in hiPSCs) +/- 3kb of STARR-seq peak centres in primed human pluripotent stem cells (PSC), mesendoderm (MES), trophoblast-like cells (TBL), mesenchymal stem cells (MSC), and neuronal progenitor cells (NPC). The majority of enhancers are not enriched for H3K27ac in any of the analysed cell types (no H3K27ac). Subsets are enriched for H3K27ac in one (Lineage-restricted) or multiple differentiated cell types (Multi-lineage). (B) Heatmap showing H3K27ac enrichment (ChIP-seq) at chromatin-repressed enhancers (mapped in hiPSCs) +/- 3kb of STARR-seq peak centres in PSC, MES, TBL, MSC, and NPC.

**Figure S4:**
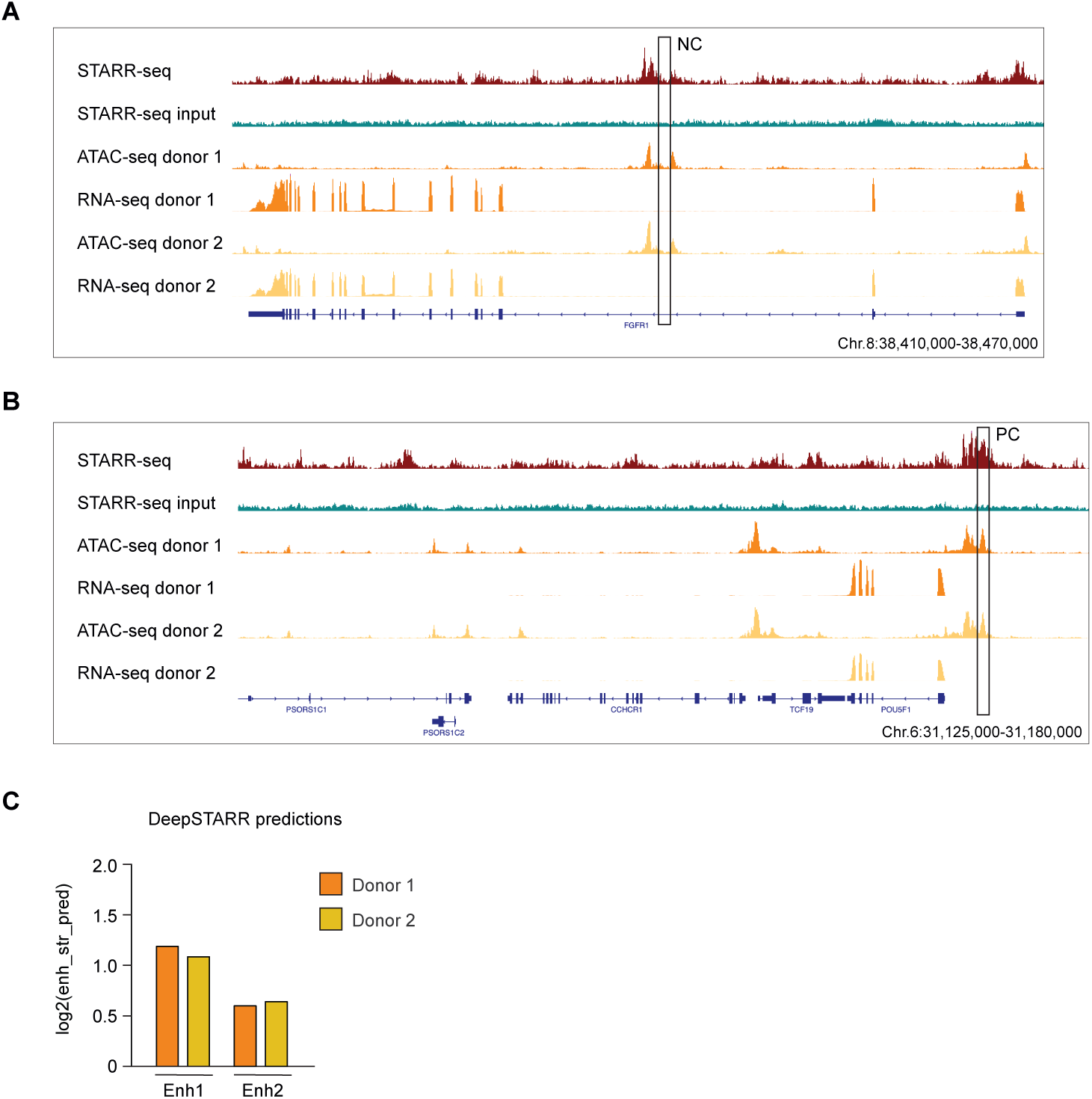
DeepSTARR predicts enhancer variant activity. (A) Coverage tracks of STARR-seq (donor 1) and ATAC-seq and RNA-seq data from two independent hiPSC lines (donor 1 and donor 2) for the negative control region used in the luciferase assay in Fig. 4C. (A) Coverage tracks of STARR-seq (donor 1) and ATAC-seq and RNA-seq data from two independent hiPSC lines (donor 1 and donor 2) for the positive control region (*POU5F1* distal enhancer) used in the luciferase assay in Fig. 4C. (C) DeepSTARR prediction for the activity of enhancer variants shown in Fig. 4C.

**Figure S5:**
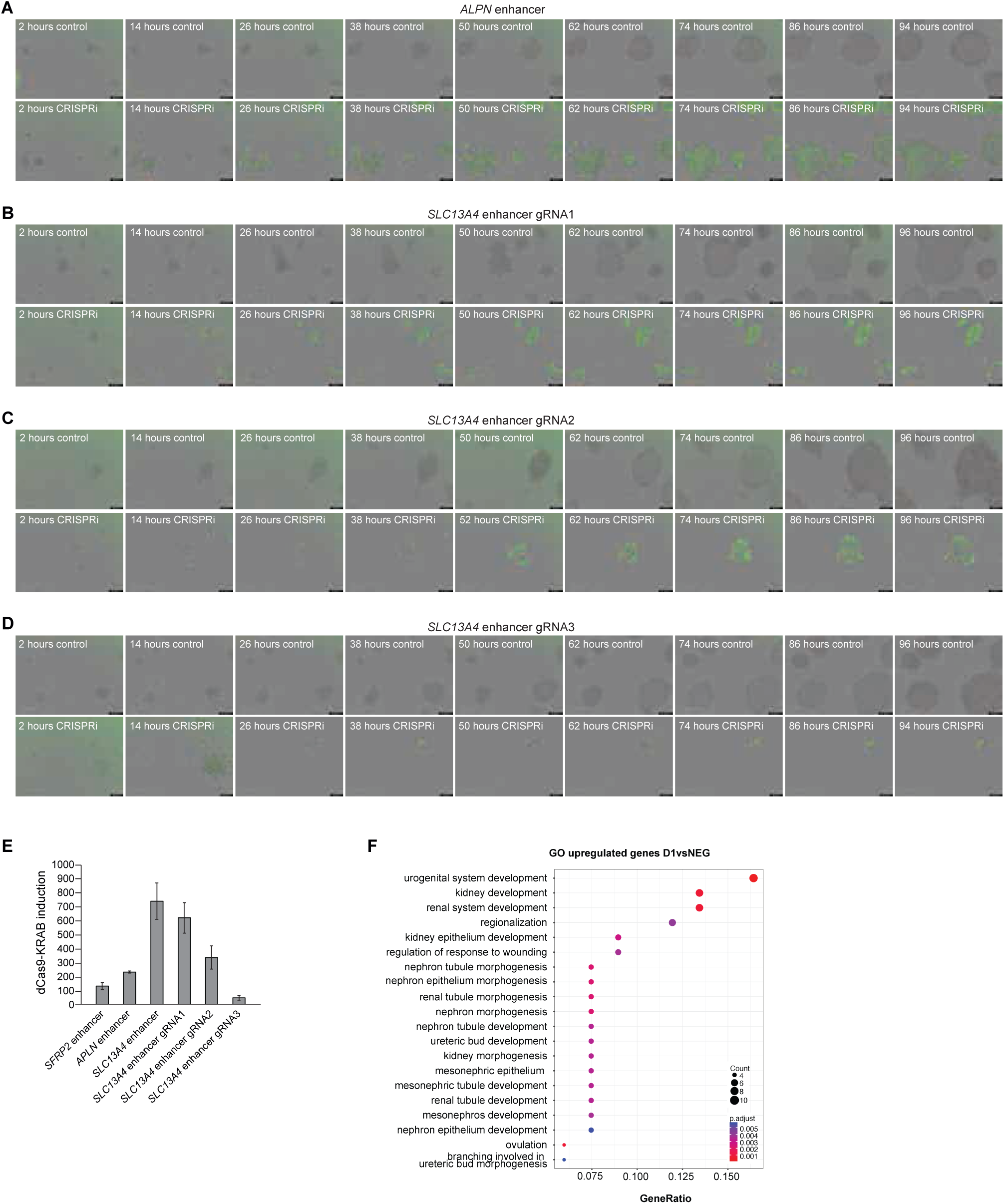
Repression of the SLC13A4 enhancer impairs cell viability. (A) Live-cell imaging of hiPSCs without (control) or with (CRISPRi) the addition of DOX and TMP to induce the expression of dCas9-KRAB, marked by GFP expression (green). Constitutive expression of three gRNAs targeting the primed enhancer of *APLN*. (B) - (D) Live-cell imaging of hiPSCs without (control) or with (CRISPRi) addition of DOX and TMP to induce the expression of dCas9-KRAB, marked by GFP expression (green). Constitutive expression of individual gRNAs (gRNA 1-3) targeting the primed enhancer of *SLC13A4*. (E) Quantitative RT-PCR for dCas9-KRAB after the addition of DOX and TMP (for 4 days) in hiPSC lines with constitutive expression of gRNAs targeting enhancers of *SFRP2*, *APLN*, or *SLC13A4*. 2^-ΔΔCt^ ± s.d.; normalisation using GAPDH (housekeeping gene) and uninduced hiPSCs as a reference (= 1). (F) GO term enrichment of differentially upregulated genes with RNA-seq datasets for hiPSCs after the addition of DOX and TMP for 1 day to induce dCas9-KRAB with constitutive expression of gRNAs targeting the enhancer of *SLC13A4* in comparison to control hiPSCs.

**Figure S6:**
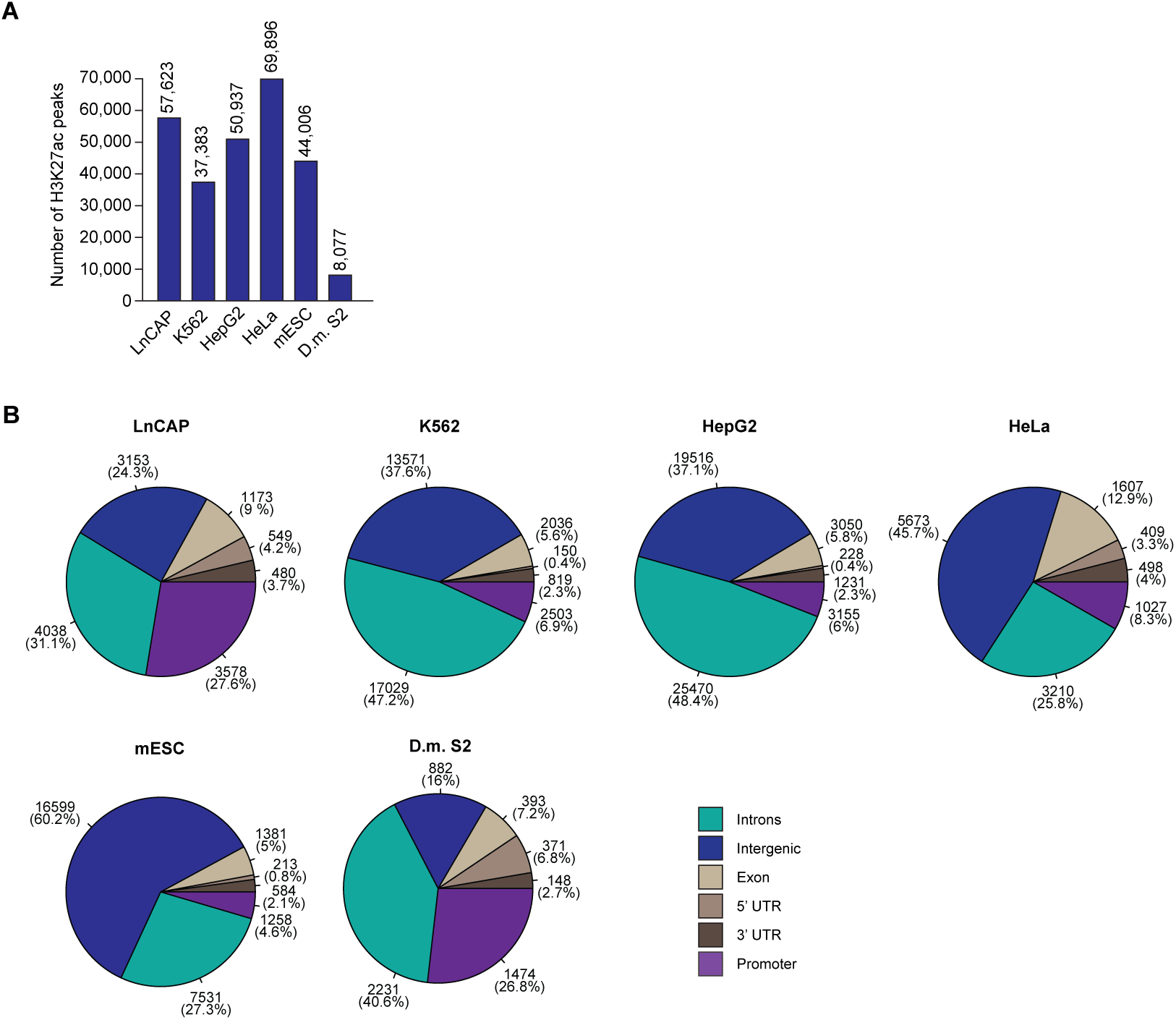
Genomic features of putative non-canonical enhancers. (A) Number of called H3K27ac peaks (ChIP-seq) for datasets described in Figure 6D. (B) Numbers (and percentage of total) of STARR-seq enhancers within indicated genomic features for datasets described in Figure 6D.

## SUPPLEMENTARY TABLES

Supplementary Table 1: Genomic coordinates of chromatin-repressed hiPSC STARR-seq enhancers

Supplementary Table 2: Genomic coordinates of active hiPSC STARR-seq enhancers

Supplementary Table 3: Genomic coordinates of primed hiPSC STARR-seq enhancers

Supplementary Table 4: Genomic coordinates of poised hiPSC STARR-seq enhancers

Supplementary Table 5: DpnII Promoter Capture Hi-C probes

Supplementary Table 6: HindIII Promoter Capture Hi-C probes

Supplementary Table 7: RT-qPCR and next generation sequencing library generation primers

Supplementary Table 8: gRNAs for CRISPRi and CRISPRa

## SUPPLEMENTARY MOVIES

Supplementary Movie 1: *SLC13A4* enhancer CRISPRi 3 gRNAs induced (hiPSC line Yoch6)

Supplementary Movie 2: *SLC13A4* enhancer CRISPRi 3 gRNAs control (hiPSC line Yoch6)

Supplementary Movie 3: *APLN* enhancer CRISPRi 3 gRNAs control (hiPSC line Yoch6)

Supplementary Movie 4: *APLN* enhancer CRISPRi 3 gRNAs induced (hiPSC line Yoch6)

Supplementary Movie 5: *SLC13A4* enhancer CRISPRi 3 gRNAs control (hiPSC line Kolf2)

Supplementary Movie 6: *SLC13A4* enhancer CRISPRi 3 gRNAs induced (hiPSC line Kolf2)

